# BCR repertoire analysis and cloning of antibody candidates targeting native and Asp7-isomerized β-amyloid

**DOI:** 10.64898/2025.12.26.696595

**Authors:** N.A. Nikolaev, M.Yu. Myshkin, E.A Kolobova, I.A. Shagina, O.I. Kechko, K. B. Varshavskaya, E. V. Barsova, T.V. Grigoreva, V.A. Mitkevich, E.M. Merzlyak, S.A. Kozin, A.A Makarov, D.M. Chudakov, S.A. Lukyanov, I.L. Grigorova, O.V. Britanova

## Abstract

Computational approaches are increasingly used to prioritize monoclonal antibody (mAB) candidates from BCR-seq datasets. However, the reliable identification of B cells encoding antibodies against rare antigenic epitopes remains challenging. We employed a repertoire-guided workflow combining antigen-tetramer sorting of B cells from immunized mice, followed by low-input bulk BCR-seq, which yielded informative clonal repertoires. Clustering analysis allowed us to prioritize IGH and IGK clonotypes potentially targeting β-amyloid and the isoAsp7 proteoform (isoD7-Aβ_1-16_), implicated in Alzheimer’s disease. We observed recurrent IGHV8-12 and IGKV1-117 usage, consistent with canonical mouse anti-Aβ responses. Focusing on isoD7-Aβ-associated candidates, we selected fourteen candidate IGH–IGK combinations for recombinant expression including variants containing IGKV8-21, IGKV8-19, IGKV2-112 together with IGHV1-75, IGHV10-1 or IGHV8-12. Several recombinant mABs showed detectable binding activity to isoD7-Aβ, including a candidate with preferential recognition of isoD7-Aβ, supporting the feasibility of the approach but underscoring the challenge of accurate chain pairing. These results provide proof of principle that repertoire-guided analysis can prioritize candidate IGH and IGK chains and propose possible heavy-light chain combinations based on their co-representation across samples while experimental testing remains necessary to identify antibodies with preferential recognition of isoD7-Aβ.

## Introduction

Alzheimer’s disease (AD) is the most prevalent form of dementia, accounting for approximately 70% of cases worldwide [1], and is pathologically defined by extracellular amyloid plaques composed of β-amyloid (Aβ) peptides and intracellular neurofibrillary tangles. Soluble Aβ oligomers, rather than insoluble deposits, are now considered the primary neurotoxic species: they disrupt synaptic transmission, impair long-term potentiation, and promote cognitive decline. Among pathogenic Aβ variants, the most common post-translational modifications are the pyroglutamate-modified Aβ (pGlu3-Aβ) and the Asp7-isomerized peptide (isoD7-Aβ). Aspartate isomerization is a non-enzymatic, age-dependent process that proceeds via a succinimide intermediate [2]. Both pGlu3-Aβ and isoD7-Aβ display an increased tendency to oligomerize, seed aggregation, and accumulate in the human brain with age [3, 4, 5]. In vitro studies demonstrate that both isoD7-Aβ and pGlu3-Aβ possess markedly greater neurotoxicity than unmodified Aβ [6, 7]. Moreover, peripheral administration of isoD7-Aβ induces plaque formation in an AD mouse model, supporting its direct role in amyloid pathology initiation in vivo [8].

Recent data on humanized IgG1 monoclonal antibodies donanemab, specific for pGlu3-Aβ (NCT04437511), and lecanemab (NCT03887455), which targets soluble aggregated species of Aβ, demonstrated a substantial reduction of brain amyloid burden and clinically modest slowing of cognitive decline in patients with early Alzheimer’s disease [9, 10]. Nevertheless, monoclonal antibodies (mAB) that specifically recognize additional abundant post-translationally modified Aβ species, such as isoD7-Aβ_1-16_, may offer benefits in Alzheimer treatment since such mAB targets pathogenic, aggregation-prone and disease-restricted Aβ forms.

In 5xFAD transgenic mice, isoD7-Aβ_1-16_-specific antibody, K11, has been shown to significantly reduce both isoD7-Aβ_1-16_ and total Aβ_1-16_ levels in the brain and to improve performance in behavioral tests [11]. These data support isoD7-Aβ_1-16_ as a promising target in Alzheimer’s disease.

The search for antigen-specific mABs typically involves immunization of a host organism followed by screening for the desired antigen-specific B cell clones. The discovery of mABs with high affinity to the antigen usually involves multiple rounds of immunization. Following immunization, B cells that are activated by antigens binding to their B cell receptors (BCRs) and receive T-cell help undergo activation, proliferation, and differentiation into short-lived plasmablasts and germinal center (GC) B cells. GCs typically form within 6-7 days and resolve within 21 days after immunization. Within these structures, B cells undergo random somatic hypermutation of their BCR genes and affinity maturation. This is a selective process where B cell clones with the highest antigen-binding affinity are progressively expanded. GC B cells can further differentiate into class-switched memory B cells and long-lived plasma cells. Secondary immunizations rapidly induce differentiation of pre-existing memory B cells into plasmablasts, as well as the recruitment of both naive and memory B cell clones into new GC reactions. Each subsequent round of GC selection following a booster immunization is believed to enhance the affinity of the antigen-specific B cell repertoire. This affinity improvement corresponds to the progressive accumulation of mutations in the complementarity-determining regions (CDRs) of the BCR heavy and light chains [12, 13,14].

In recent years, together with classical hybridoma technology, other strategies have been developed to generate recombinant mAB directly from peripheral blood. It includes techniques based on single-cell sequencing or sorting of antigen-binding B cells, immortalization and screening of B cells with Epstein–Barr virus (EBV) [15–18]. The EBV transformation approach was successfully applied to generate neutralizing mABs against SARS-CoV [19], although it required polyclonal activation and labor-intensive screening to isolate high-affinity B cell clones.

Currently the growing availability of high-throughput B cell receptor (BCR) repertoire datasets has driven the development of computational tools for prediction antigen-associated BCRs. A recent study introduced algorithms capable of clustering CDR sequences to identify epitope-specific BCRs [20]. In parallel, language model-based BCR embeddings have demonstrated improved performance in predicting receptor specificity [21].

The primary objective of this study was to develop and explore a bioinformatics workflow for the analysis of antigen-specific B cell repertoires and prioritization of monoclonal antibody candidates following immunization of BALB/c mice with isoD7-Aβ□ -□□. Using a previously established immunization strategy [22], repertoire-derived antibody candidates were selected computationally and subsequently evaluated experimentally as recombinant mABs for binding to isoD7-Aβ and non-isomerized Aβ. This study was designed as a proof-of-principle assessment of the feasibility of repertoire-guided candidate selection rather than as a validation of a predictive antibody-discovery pipeline.

## 2. Results

### 2.1. Isolation of enriched antigen-specific B cells after isoD7-Aβ-targeting immunization in BALB/c mice

To promote isoD7-Aβ_1-16_-specific B cell response, BALB/с mice were intraperitoneally immunized with ovalbumin (OVA) conjugated to isoD7-Aβ_1-16_ peptide via C-terminus (with peptide N-terminus facing outside) in Ribi adjuvant (**Figure 1A**). The immunization induces B cell response to isoD7-Aβ_1-16_, including B cells cross-reactive and not cross-reactive to Aβ - [22].

**Figure 1.**
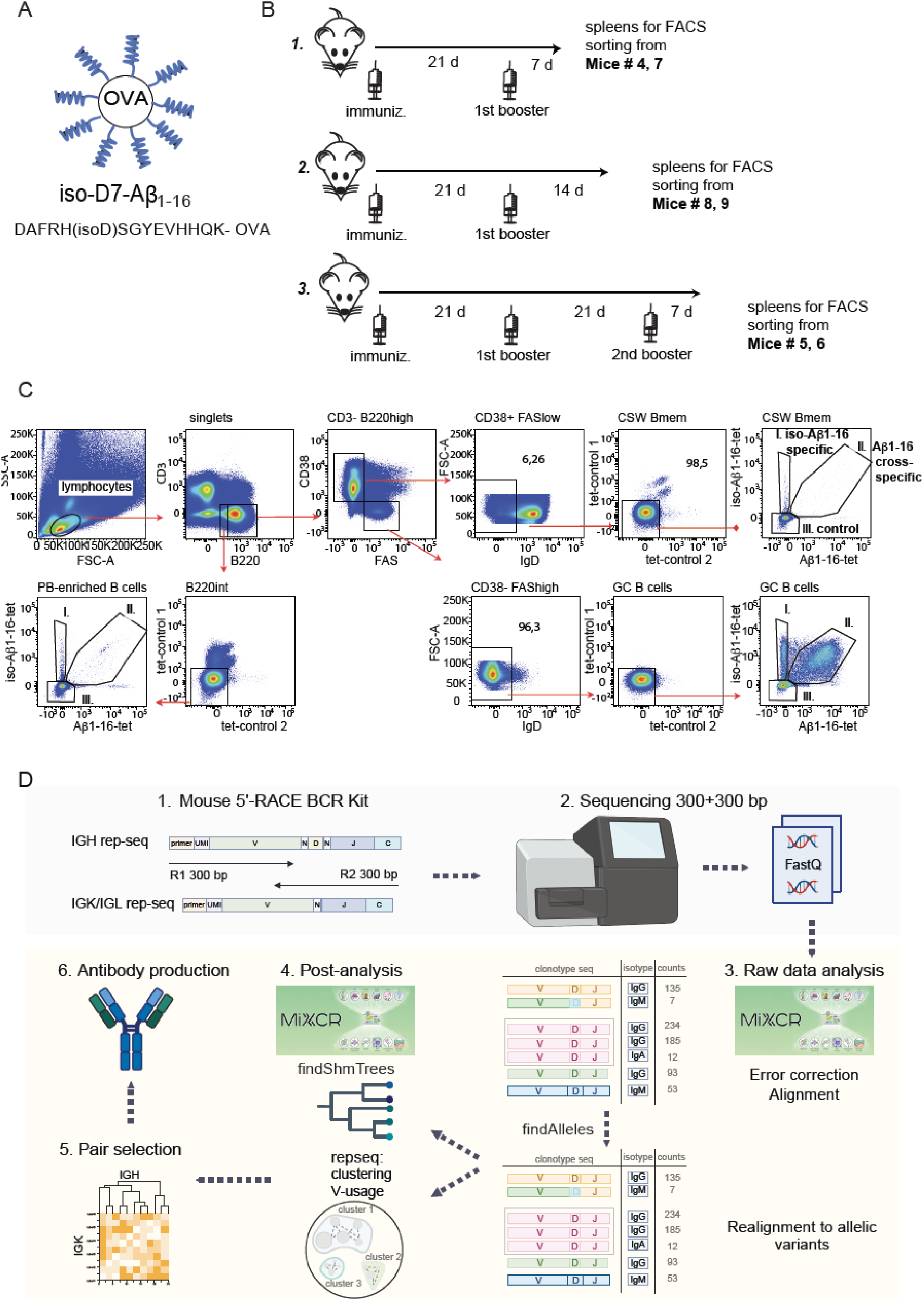
Experimental strategy for FACS sorting Aβ_1-16_- and/or isoD7-Aβ_1-16_-specific and non- specific subpopulations of B cells. (A, B) Immunization strategy. BALB/c mice were immunized intraperitoneally and subsequently boosted with OVA-isoD7-Aβ_1-16_ conjugate antigen (A) in the Ribi adjuvant according to the schedule shown in (B) (groups 1-3). Spleens were collected at the indicated time points for immunofluorescent staining with a mixture of fluorochrome-conjugated antibodies against cell surface markers and fluorescent tetramers with isoD7-Aβ_1-16_antigen, Aβ_1-16_ antigen and two decoy tetramers (tet-controls 1 and 2). (C) Gating strategy enables analysis of CD3^-^B220^high^CD38^+^ FASlow IgD^-^ B cells enriched with class-switched memory B cells CSW Bmem, CD3^-^B220^high^CD38^-^ FAS^high^IgD^-^ germinal center B cells and CD3^-^B220^int^ cells enriched for plasmablasts. These populations were further subdivided into specific to iso-D7-Aβ_1-16_ (gate I), cross-specific to iso-D7- Aβ_1-16_ and Aβ_1-16_ (gate II) or non-specific or specifically non-binding to the tetrameric antigens (gate III). (D) Workflow for BCR repertoire analysis and selection of antibody candidates. 5′-RACE–based mouse BCR repertoire sequencing: IGH and IGK/IGL libraries are generated with UMI-containing primers and sequenced using 2 × 300 bp paired-end reads. Raw sequencing data are processed with MiXCR for read alignment, UMI-based error correction, clonotype assembly and isotype annotation. Post-analysis includes identification of allelic variants (findAlleles) and reconstruction of somatic hypermutation trees (findShmTrees), V-usage analysis. Clonotypes are clustered at the amino-acid level (repseq.clustering) to identify IGH–IGK pairs enriched in antigen-specific B-cell samples. Selected IGH/IGK combinations are expressed as recombinant mABs and tested functionally. Created in https://BioRender.com

Based on this knowledge, three immunization schemes have been selected to collect B cells with progressive accumulation of clones with improved BCR affinity to the immunization antigens (Figure 1B). All mice immunized with OVA-isoD7-Aβ_1-16_ in Ribi were boosted with the same antigen/adjuvant combination in 21 days. Spleens were collected from the boosted mice in 7 days (group 1) and in 14 days (group 2). Spleens were also collected from mice at 7 days after they received a 2nd booster immunization (group 3). It is expected that GC B cells in groups 1, 2, and 3 have progressively improved their affinity to the antigens. The mice in the 3rd group were also expected to accumulate some short-living plasmablasts from high-affinity memory B cells generated before the second boost. To identify Aβ -specific germinal center (B220 CD3 CD38 FAS^high^IgD) and B cells enriched for class-switched memory cells, splenocytes were stained with fluorochrome-conjugated antibodies against the indicated surface markers and tetramers composed of the fluorescent streptavidins conjugated with either biotinylated isoD7-Aβ_1-16_ or Aβ_1-16_ peptides (**Figure 1C).** As shown previously [22], cross-reactive B cells bind both tetramers simultaneously, whereas B cells predominantly specific for isoD7-Aβ_1-16_ bind only the isoD7-Aβ_1-16_ tetramer.

Based on this gating strategy, we have sorted isoD7-Aβ_1-16_-specific GC B cells (iso-specific) and B220 CD3 CD38 FAS IgD B cells enriched for class-switched (CSW) Bmem (40 to 400 and from 34 to 213 cells per sample, correspondingly) for BCR profiling from the indicated groups of mice (**Figure 1C, gate I, Table S1**). As controls, we also sorted GC and memory B cells cross-reactive with isoD7-Aβ_1-16_and Aβ_1-16_ (cross-specific) (**Figure 1C, gate II, Table S1**) with yields ranging from 24 to 400 and from 8 to 65 cells per sample correspondingly, Additionally, non-specific GC B cells (non-binding either isoD7-Aβ_1-16_ or Aβ_1-16_) were collected at 1000 cells per sample in replicates (**Figure 1C, gate III**). In group 3, we additionally observed an accumulation of B220 int/low B cells binding the isoD7-Aβ_1-16_ and Aβ_1-16_ tetramers (**Figure 1C**). These cells are likely to be early plasmablasts that have already downregulated B220 but still have some BCRs on the surface before they are completely downregulated in more mature plasma cells. One of the mice (#6, **Figure 1B**, group 3) demonstrated the strongest response and allowed us to sort two replicas by 400 isoD7-Aβ_1-16_-positive CD3^-^B220^int^ B cells (**Table S1**).

### 2.2. BCR repertoire analysis prioritized candidate IGH and IGK clonotypes associated with isoD7-Aβ1-16-specific B-cell populations

BCR clonotypes were extracted with MiXCR using UMI-aware assembly for error correction and high-confidence clonotype calling (**Table S2**). The overall workflow is shown in **Figure 1D**. On average, sequencing depth was sufficient for reliable error correction with >30 reads per UMI. The number of UMI-tagged IGH cDNA molecules per input cell showed substantial variability across samples (**Figure S1**). This variability likely reflects both technical factors (e.g., cell lysis efficiency, cDNA synthesis) and biological heterogeneity, including differences in the differentiation states of B cell populations across samples. Naïve B cells are known to express significantly lower levels of BCR transcripts compared to antibody-secreting B cells [23]. As a result, samples containing higher proportions of activated or more differentiated B cells are expected to yield greater numbers of BCR transcripts per input cell.

To identify candidate BCR clonotypes associated with iso-specific B cells, we employed several strategies:

1. Clustering analysis to group similar IGH and IGK amino acid sequences and define putative antigen-specific clonotypes
2. Analysis of enriched IGHV and IGKV rearrangements in repertoires of iso-specific and cross-specific samples
3. Somatic hypermutation (SHM) tree reconstruction using MiXCR to identify lineages undergoing somatic hypermutation.

In this study, we analyzed “clonotype groups”, which we defined as a unique combination of V-segment, J-segment, and amino acid CDR3 sequence (CDR3aa), rather than using MiXCR-defined clonotypes directly. This approach was intended to increase sensitivity while preserving features most relevant to antibody binding, which is primarily determined by the V-segment type and CDR3 sequence. We extracted all unique V-J-CDR3aa combinations from the entire dataset and assigned them continuous numbering. Throughout most of the analysis (except the final full-sequence extraction for *in vitro* tests), we applied a “pooling” procedure that merged clonotype-group records into a single clonotype within each repertoire thereby ignoring substitutions in the CDR1/2 and FR1/2/3 regions.

We first analyzed IGHV gene-segment usage which had cumulative frequency > 0.05 in at least one repertoire sample (**Figure S2**). We found that IGHV10-1 and IGHV14-3 were significantly enriched in sorted isoD7-Aβ_1-16_-specific B cell samples, while IGHV1-69, IGHV3-1, and IGHV8-12 showed a trend toward enrichment (**Figure 2B**), (see Material and Methods). Using a filter adjusted for the representation of IGHV10-1 and IGHV14-3 in the isoD7-Aβ_1-16_-specific (iso-specific) repertoires, six clusters were selected (**Figure 2C, Figure S3**). For instance, cluster 1 comprised clonotypes for which 95% of the total UMI count was derived from iso-specific clonotypes. Cluster 1 was the most widely shared across mice and consisted of IGHV10-1 clonotypes from mice #4, 5, 6, 7 and 9 and contained fourteen iso-specific nodes out of sixteen total nodes. In addition, cluster #56 was notable in that 73% of its total UMIs were contributed by isoD7-Aβ_1-16_-specific IGHV10-1 clonotypes from mice #4 and #6. This cluster displayed a highly similar CDR3 consensus to cluster 1, but used a different IGHJ gene segment (**Figure 2C, Table S3**). Another promising cluster, #18, comprised IGHV14-3 clonotypes and met both selection criteria: by specificity and IGHV usage. Next, we selected twelve clusters that consisted only of clonotypes originating from isoD7-Aβ_1-16_-specific sorted B cells. Additionally, we built a multiple sequence alignment (MSA) for full-length nucleotide sequences of all functional IGH clonotypes, computed from it a distance matrix of pairwise Hamming distances, and embedded the sequences using multidimensional scaling (MDS). This analysis revealed several dense clusters that were associated with IGHV segment usage and enriched for clonotypes from the sorted B cell populations with either iso-specific or cross-specific reactivity (**Figure 2D**).

**Figure 2.**
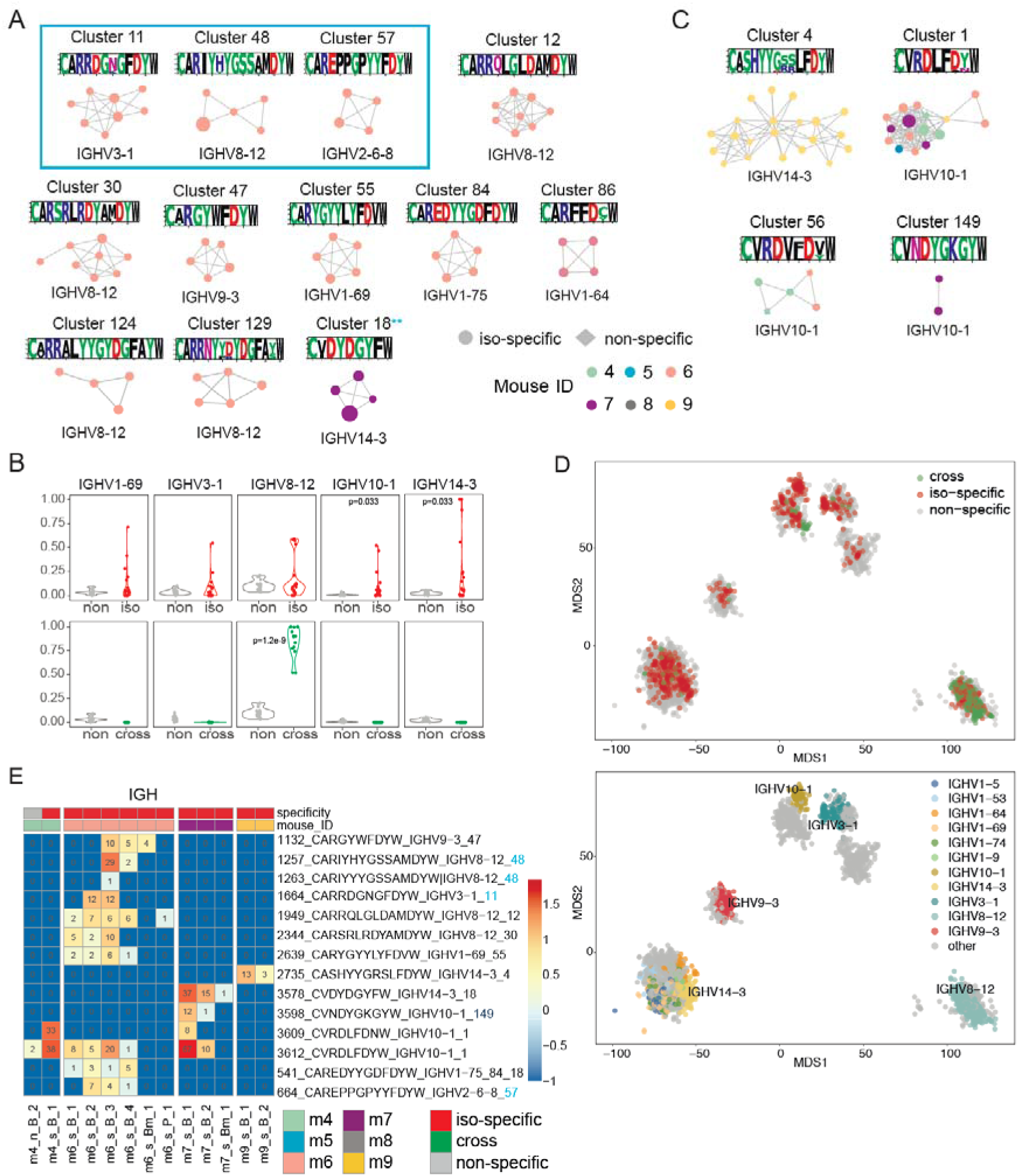
Repertoire analysis of IGH sequences in sorted B cell populations specific for isoD7-Aβ_1-16._ (A) Representative examples of IGH clonotype clusters shown as network graphs built from repertoires of sorted B cell populations. Clonotypes were grouped if clonotypes used the same IGHV gene and differed by no more than one amino acid in the CDR3 region (≤1 aa mismatch). Shown clusters met the following criteria: contained only nodes derived from sorted isoD7-Aβ1-16-specific B cell samples (≥4 nodes) and the cumulative UMI count of nodes within each cluster was ≥10. Colors indicate the mouse of origin (m4–m9). Blue box highlights clusters selected for further analysis, two stars indicate cluster #18 met both filtering criteria. Consensuses and UMI count in iso-specific repertoires are shown above each cluster. (B) The top five IGHV segments were selected using a mean frequency threshold of >0.035 across all samples. IGHV abundances were compared between repertoires of non-specific (N = 20) and isoD7- Aβ_1-16_-specific (N = 20) as well as between non-specific (N = 20) and cross-specific (N = 13) repertoires from sorted B cells. P values were calculated using one-sided t-test and adjusted for multiple comparisons using the Benjamini–Hochberg false discovery rate (FDR) correction. (С) Shown are the top iso-specific clusters with cumulative UMI counts >100 that incorporated IGHV10-1 and IGHV14-3 and contained at least two nodes derived from sorted isoD7-Aβ_1-16_-specific B cell samples. Colors indicate the mouse of origin (m4–m9). (D) Global embedding of IGH repertoires generated by multidimensional scaling (MDS) based on pairwise Hamming distances between aligned full-length nucleotide sequences. Each dot represents one clonotype, and dot size is proportional to its UMI count in the repertoire. Dots are colored by specificity of sorted B cell samples, V-gene segment usage. (E) Heatmaps showing the abundance of selected IGH clonotype groups across samples, expressed as log10-transformed UMI counts. Samples are ordered by specificity (iso-specific, cross- specific, or non-specific) and by mouse ID. UMI counts in the sample are shown in each box. Blue color shows clonotype groups selected for recombinant expression.

In parallel, we reconstructed possible lineage trees based on SHM patterns to identify IGH clonotypes likely undergoing antigen-driven selection. Interestingly, a substantial portion of SHM tree sequences retained the IgM isotype, which is consistent with early-stage responses and may also be compatible with extrafollicular diversification; however, these possibilities cannot be distinguished from the present data. No SHM trees meeting the selection criteria were found in mice #7, #8, or #9. A key limitation of SHM lineage reconstruction in this study is the relatively low sequencing depth, which may restrict the ability to detect expanded B cell lineages. Among selected clusters, clonotypes from cluster #11 (IgM) in mouse #6 was incorporated into SHM lineage tree #170 and clonotypes from cluster #84 (IgM and IgG) contributed to SHM tree #2 (**Figure S4**). SHM trees were also detected in non-specific repertoires, but these were interpreted as background clonal diversification unrelated to the iso-specific response.

Taken together, these two filtering steps allowed us to prioritize clonotype groups from selected clusters which were abundant in repertoires of isoD7-Aβ_1-16_-specific B cells.

We focused on the abundant clonotype groups (**Figure 2E**) which were detected in at least two isoD7-Aβ_1-16_-specific B samples and had a UMI count of at least 10 isoD7-Aβ_1-16_-specific samples (**Figure 2E**). In total, seven IGH clonotype groups from clusters #1, 11, 18, 48, 57, 84 were included in the subsequent analysis across mice #4, 6, 7. Four of these clonotype groups used IGHV14-3, IGHV10-1 or IGHV8-12.

Next, we performed clustering analysis of the immunoglobulin light chain repertoires. Although both kappa-(IGK) and lambda-chain (IGL) clonotypes were recovered, IGL clonotypes were extremely rare in this dataset. IGL clonotypes were detected in only three samples. We therefore restricted downstream light-chain clustering and candidate selection to IGK repertoires.

IGK clusters displayed substantial promiscuity and also included clonotypes originating from non-specific and cross-specific B cell subsets across different mice. Analysis of the IGKV usage nevertheless revealed significant enrichment of IGKV1-110, IGKV1-117, IGKV8-19 in iso-specific samples compared to non-specific ones (**Figure S2 and S6**).

We therefore focused on clusters containing clonotypes in repertoires of sorted isoD7-Aβ_1-16_-specific B cells (**Figure 3A**) and ranked them by the cumulative UMIs of the clonotypes from isoD7-Aβ_1-16_-specific samples (**Table S4**). Additionally, we embedded full nucleotide sequences of functional IGK clonotypes using MSA and MDS as described above for IGH repertoires (**Figure 3C, Figure S5**). This analysis revealed several dense regions enriched in IGK clonotypes from iso-specific samples, particularly those using IGKV1-110, IGKV1-117, IGKV8-19, IGKV8-21, and IGKV6-15 (DBSCAN clusters, **Figure S5**), in agreement with the IGKV usage and iso-specific cluster analysis.

**Figure 3.**
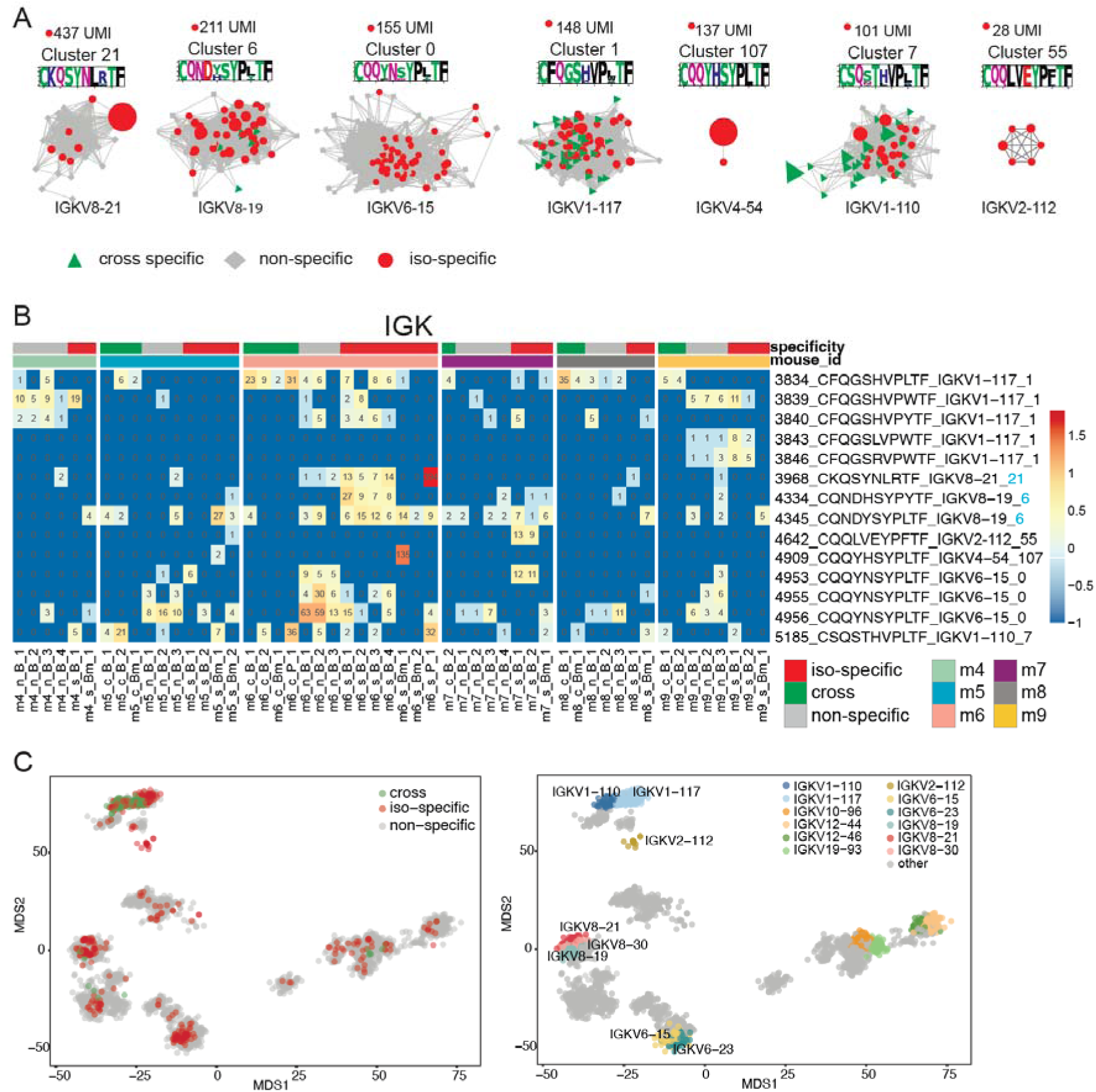
Repertoire analysis of IGK sequences in sorted B cell populations specific to isoD7-Aβ_1-16_. (A) Representative examples of IGK clonotype clusters shown as network graphs constructed from the repertoires. Each node represents a unique clonotype with edges showing sequence similarity (with 1 mismatch in CDR3 amino acid). Clonotypes from isoD7-Aβ_1-16_ sorted B cells are shown as circles, and cross-reactive clonotypes as triangles. (B) Heatmaps showing the abundance of selected IGK clonotype groups across samples within clusters #0, #1, #6, #7, #21, #55, #107 expressed as log10- transformed UMI frequencies. Samples are ordered by specificity (iso-specific, cross-specific, or non-specific) and by mouse ID. UMI counts in each sample are shown in the corresponding box. (C) Global embedding of IGK repertoires generated by classical multidimensional scaling (MDS) based on pairwise Hamming distances between aligned full-length nucleotide sequences. Each dot represents one clonotype, and dot size is proportional to its UMI count in the repertoire. Dots are colored by specificity of sorted B cell samples, IGKV gene segment usage.

For functional testing, we prioritized IGK clusters using two complementary criteria: high representation in isoD7-Aβ_1–16_-specific repertoires and enrichment for antigen-specific clonotypes. We selected six IGK clusters (#0, 1, 6, 7, 21, 107) with the UMI count exceeding 100 (**Figure 3A, Table S4**). Although cluster 0 contained a relatively low proportion of isoD7-Aβ_1-16_-specific clonotypes, accounting for only 15% of the total cluster UMI count, we still considered it a candidate public IGK cluster that might contribute to isoD7-Aβ_1-16_-specific antibodies. For functional testing, we selected IGKV8-21 and IGKV8-19 clonotypes belonging to the most iso-specific enriched clusters, 21 and 6, respectively. Notably, antibody K11, previously identified as highly specific for isoD7-Aβ_1-16_, used the IGKV8-21 [11]. In addition, we identified two small IGK clusters (#107 and #55) composed exclusively of isoD7-Aβ_1-16_-specific clonotypes. For further analysis, we selected cluster #55, as it contained more than two nodes and was reproducibly detected in two independent replicates from mouse #7 (**Figure 3B**).

Based on co-representation across isoD7-Aβ_1-16_-specific samples and mice, together with cluster enrichment and sequence-similarity criteria, we proposed several candidate IGH–IGK combinations for recombinant production (**Table 1**). Pairing was assigned to clonotype groups that were recurrently detected in the same mice belonging to selected IGH and IGK clusters. In total, seven IGH clonotypes representing six IGH clusters (#1, #11, #18, #48, #57, and #84) and five IGK clonotypes representing five IGK clusters (#0, #1, #6, #21, and #55) were selected for recombinant antibody production and assembled into 14 IGH–IGK combinations (Table 1). Thus, IGHV3-1 clonotype #1664 derived from cluster #11 could be paired with IGK clonotype #4345 and #3968, both of which were detected in samples of sorted isoD7-Aβ_1-16_-specific B cells from mouse #6 (**Figure 3B**).

**Table 1.**
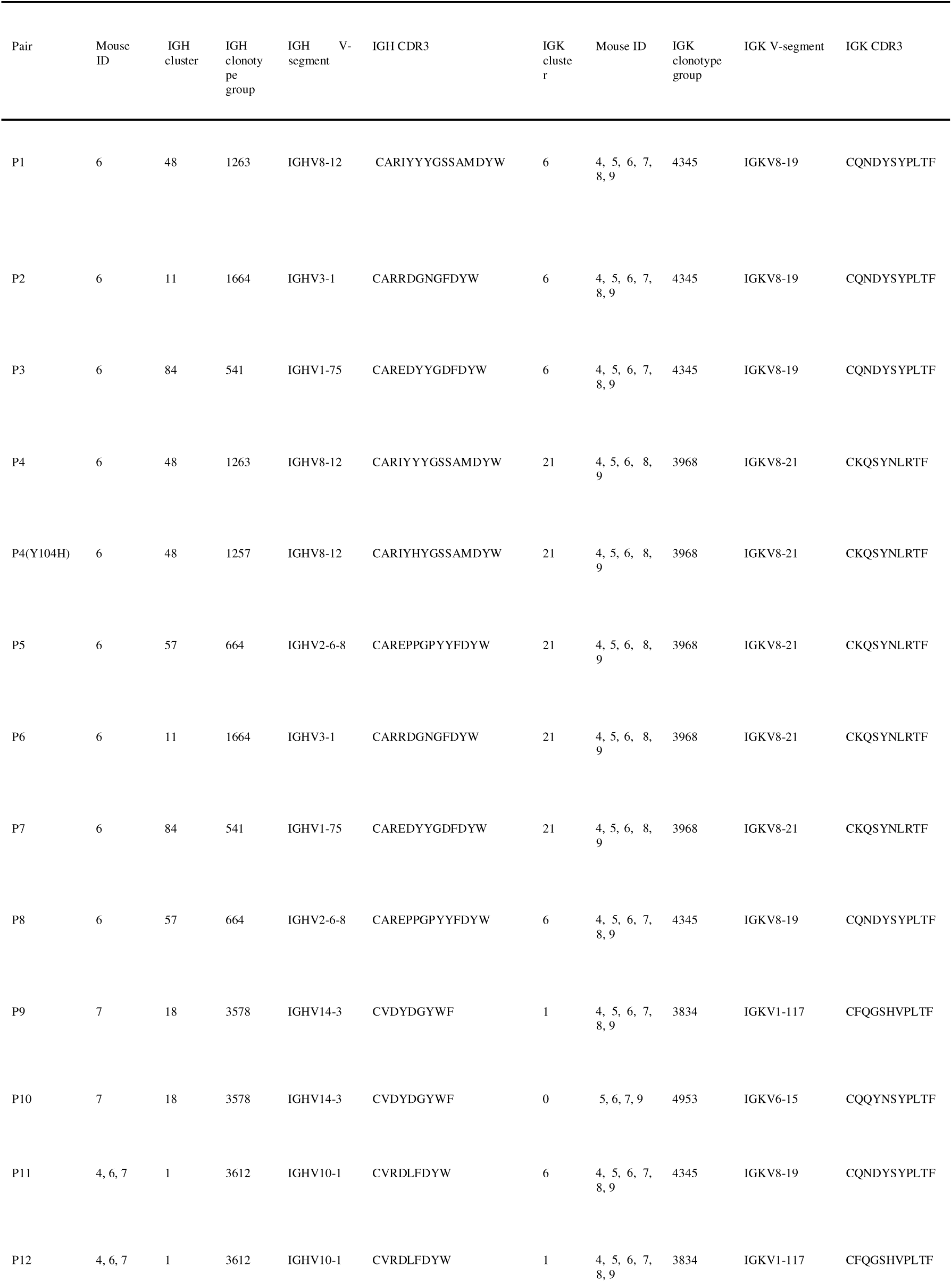
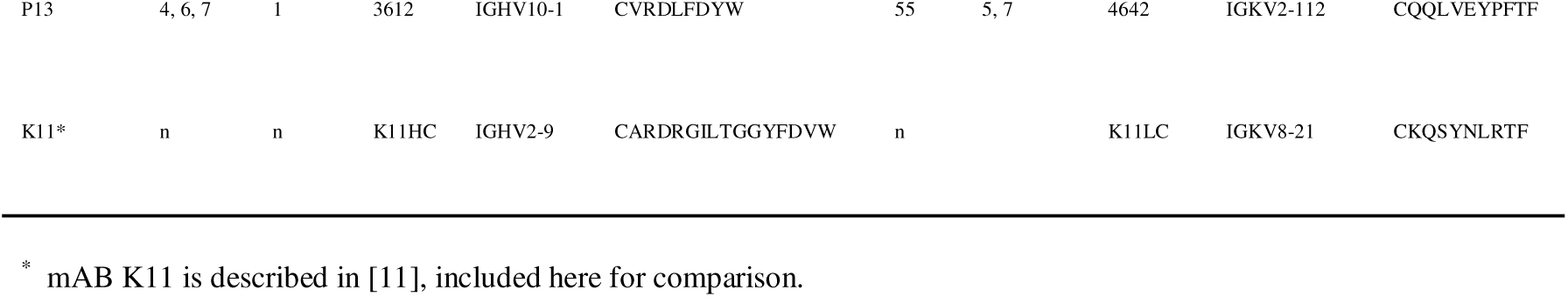
Description of the selected IGH and IGK chains for pairing and production of recombinant mABs.

In addition, cluster #48 was predominantly enriched in IGM clonotypes and included clonotype #1257 (IGHV8-12–IGHJ4; CDR3, CARIYHYGSSAMDYW). A closely related clonotype, #1263, used the same IGHV8-12 and IGHJ4 segments and differed by a single amino acid in the CDR3 region (CARIYYYGSSAMDYW; H ->Y), but was detected as an IGHG1 clonotype. For recombinant expression, we selected both #1257 and #1263. Сonotype #1263 was included to determine whether class-switched variant within the same convergent cluster differed in antigen recognition. These IGH clonotypes might be paired with the two abundant IGK clonotypes #4345 and #3968 (**Figures 2 and 3**).

Taken together, pairing analysis based on enrichment in isoD7-Aβ1–16-specific repertoires, recurrence across independent samples and mice, BCR sequence clustering, and SHM lineage information was used to select heavy-and light-chain combinations for recombinant antibody production and subsequent functional validation (Table 1).

### 2.3. Identification of shared IGHV and IGKV segments previously described as recognizing Aβ

In parallel, we asked which IGH and IGK clonotypes might be Aβ-specific, enabling comparison with previously published studies. Using the same filtering strategy, we selected fourteen IGH clusters from B cells sorted as isoD7-Aβ_1-16_- and Aβ-specific (cross-specific) (**Figure S7**). Clonotypes in thirteen clusters used IGHV8-12 and one used IGHV8-8 (**Figure S7, Table S3**). Only four clusters contained more than three nodes. IGHV8-12 segment was the only significantly enriched among IGH compared to non-specific samples (**Figure 2B**). IGKV1-110 and IGKV1-117 segments were significantly more represented among IGK in cross-specific samples relative to non-specific ones (**Figure S6)**.

Notably, IGKV1-110 and IGKV1-117-containing clusters dominated the cross-specific repertoire, with UMI counts of 205 and 194, accounting for 54% and 41% of total cross-specific UMIs, respectively (**Table S4)**. The most abundant cross-specific clonotype groups are shown in **Figure S8**. The IGHV8-12 and IGKV1-110 segments have been previously reported to assemble N-terminal anti- Aβ mABs in immunization of mice [24]. The identified HCDR3s did not exactly match previously reported anti-Aβ antibodies, consistent with published data showing substantial variability of IGH CDR3 among murine anti-Aβ mABs. Nevertheless, several clonotypes displayed a certain pattern, including Tyr and Asp in CDR3, consistent with an IGHV8-biased anti-Aβ-response (**Figure S9**).

### 2.4. Recombinant expression and binding validation of repertoire-derived mABs to isoD7-Aβ

Using seven IGH and five IGK chains selected in the repertoire analysis, we assembled and expressed fourteen recombinant mAB variants (P1-P13 + P4(Y104H), differing from P4 by a single amino-acid substitution in heavy chain CDR3) with human IgG4 isotype (**Table 1**). First, we tested specific activity against isoD7-Aβ_1-16_ and Aβ_1-16_ coupled with BSA in ELISA. The isoD7-Aβ_1-16_-specific antibody (K11) previously published in [11, 25] was used as positive control. The variable regions of K11 are encoded by IGHV 2-9 and IGKV 8-21 segments. For comparative analysis, K11 was produced in the same format as human IgG4 isotype. In ELISA, the mAB variants P3 (IGHV1-75 with IGKV8-19), P4 and P4(Y104H) (IGHV8-12 with IGKV8-21), and P13 (IGHV10-1 with IGKV2-112), showed activity against both isoD7-Aβ_1-16_ and Aβ_1-16_, albeit weaker than that of K11 (**Figure 4A, B**). Identified IGK in P4 belonged to IGKV8-21 and differed by only four amino-acid substitutions including one in CDR1 and the remaining three in framework regions, from K11 IGK. Notably, the ELISA readout did not discriminate between modified and unmodified BSA-Aβ_1-16_ peptides for all mABs except P4(Y104H) and P13, perhaps due to the multimeric peptide presentation in complex with BSA that increases effective avidity of mABs binding to the immobilized antigens thus reducing the discrimination of mAB antigen-binding site affinity to Aβ_1-16_ versus isoD7-Aβ_1-16_. As summarized in Table 1, several candidates shared identical IGH (i.e., P1/P4 with HC #1263), yet differed in IGK pairing. The fact that several pairings (*e.g*., IGHV10-1 HC #3612 with IGKV2-112 LC #4642) produced binding, while alternative pairings with the same IGH or IGK did not (*e.g*., IGHV10-1 HC #3612 with IGKV8-19 LC #4345 or IGKV1-117 LC #3834), suggests that pairing rather than V-gene usage alone drives functional specificity.

**Figure 4.**
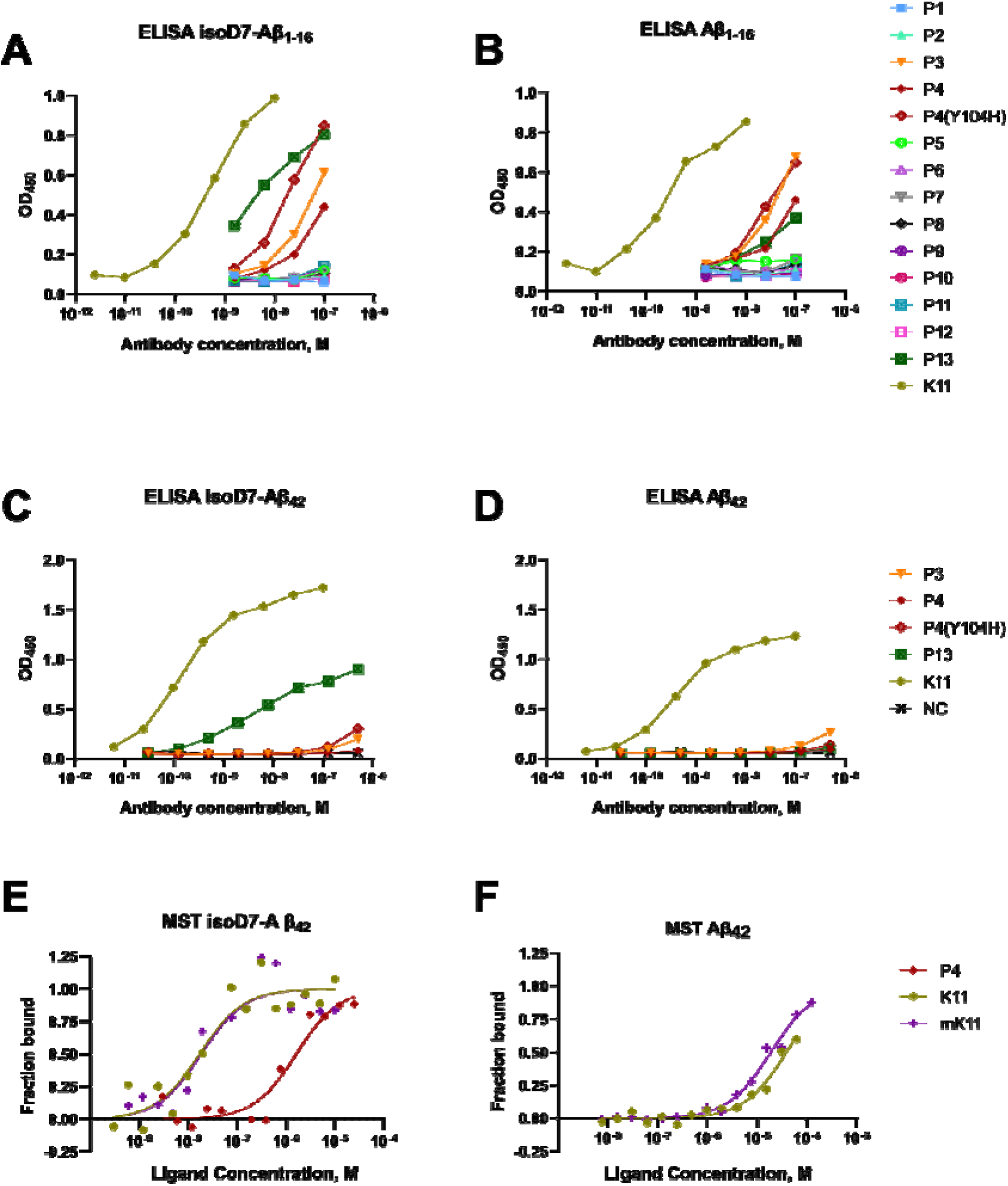
Evaluation of candidate mABs for binding to native and isoD7-modified Aβ peptides using fragment of isoD7-Aβ or Aβ, assessed by ELISA. K11 is shown as a positive control. mABs P3, P4, P4(Y104H) and P13 show binding with both antigen isoforms, albeit weaker than that of K11. Each experimental point was performed in two technical replicates. (C–D) Binding of mABs P3, P4, P4(Y104H) and P13 with full-length isoD7-Aβ_42_ or Aβ_42_, assessed by ELISA. K11 is shown as a positive control; NC – non-specific human antibody control. P3, P4(Y104H) and P13 bind isoD7-Aβ_42_, and P3 binds Aβ_42_, in all cases weaker than K11. Each experimental point was performed in two technical replicates. (E–F) Representative MST binding curves of mAB P4 with isoD7-Aβ_42_, and K11 (positive control) and mK11 (IGH from K11 paired with IGK from our data) with isoD7-Aβ_42_ or Aβ_42_. K11 and mK11 demonstrate comparable binding, while P4 binds isoD7-Aβ42 ∼100 times weaker than K11. No binding was detected for P4 with Aβ_42_, and for P3, P4(Y104H) and P13 with either isoD7-Aβ_42_or Aβ_42_ (data not shown).

Then we used antibodies P3, P4, P4(Y104H) and P13, which showed binding with BSA-conjugated N-terminal region of isoD7-Aβ and Aβ, in ELISA with full-length isoD7-Aβ_1-42_ or Aβ_1-42_ (Aβ42) (**Figure 4C, D**). K11 was used as positive control, and, unlike with N-terminal regions, bound isoD7- Aβ42 significantly stronger than Aβ42. P4 showed almost no binding with both isoforms, P3 bound both weakly. P4(Y104H) showed weak isoD7-Aβ42 binding while P13 displayed more pronounced activity. Notably, unlike K11, neither P4(Y104H) nor P13 showed detectable binding to Aβ42, suggesting a preference for isoD7-Aβ42. It was not possible to estimate any of the studied antibodies’ affinities from this experiment.

Additionally we performed an ELISA in which the concentrations of isoD7-Aβ42 and Aβ42 were varied (Figure S10, Supplemental Methods). K11 showed a clear preference for isoD7-Aβ42. P4 showed only a small difference between the two peptides, whereas P4(Y104H) consistently produced a higher signal with isoD7-Aβ42 than with Aβ42, albeit the signal was consistently weak for both of them. The strongest discrimination among the repertoire-derived mAbs was observed for P13, which showed a concentration-dependent increase in binding to isoD7-Aβ42, while the signal for Aβ42 remained close to background. Some differences in apparent binding between the two ELISA experiments may reflect differences in assay format and in the accessibility of the N-terminal epitope. Overall, these results further support preferential recognition of isoD7-Aβ42 by P13.

Finally, we attempted to validate the four candidate mABs’ binding in microscale thermophoresis (MST) using fluorescently labeled mABs. In MST, a localized temperature gradient generated by an infrared laser causes molecules to move, and the measured fluorescence signal in the heated spot is altered, while ligand interaction can affect thermophoretic mobility. Since MST is performed in solution without immobilization and wash steps, it can yield more accurate d estimation compared to plate-based assays for weak or conformation-sensitive interactions [26].

Using MST, we compared the binding affinity of the four candidate mABs to isoD7-Aβ42 and Aβ42 peptides with that one for K11. We also included mK11, a hybrid mAB, in which the K11 IGH was paired with the P4 (IGKV8-21) IGK chain. K11 and mK11 exhibited comparable Kd values when binding isoD7-Aβ42 (Kd = 18.9 ± 6.8 nМ for K11; 19.8 ± 7.7 nМ for mK11, presented here as mean ± SD) (**Figure 4E**) and also comparable, roughly 500-1000-fold weaker binding with Aβ42 (Kd 20 μM for K11; Kd 10 μM for mK11) (**Figure 4F**). P4 showed roughly 100-fold lower affinity to isoD7-Aβ42 (1.6 ± 0.2 μM) (**Figure 4E**) and no detectable binding with Aβ42. P3, and, less expectedly, also P4(Y104H) and P13 displayed no detectable binding within the tested concentration range of both isoforms, which may reflect assay-specific effects, including possible sensitivity to fluorescent labeling. Assay results for best mABs tested are summarized in **Table S5**.

Together, the ELISA results for P13 and P4(Y104H), as well as MST results for P4, indicate that the repertoire-guided approach identified antibodies with preferential recognition of isoD7, although our results also highlight the relatively low hit rate of the current repertoire-based prediction and demonstrate that correct heavy–light chain pairing is essential for recovering functional isoD7-Aβ- specific mABs.

## 3. Discussion

High-throughput BCR-seq of memory B and plasma cells is widely used to recover monoclonal antibody candidates from immune repertoires. However, recombinant production and functional screening of all potentially relevant clonotypes are not yet scalable for routine implementation. Several approaches have been developed to examine or predict BCR specificity *in silico*, such as clustering-based methods [20] and language model-based frameworks [21]. Repertoire-guided bioinformatics approaches can facilitate prioritization of high-confidence candidates for production and further validation, thereby reducing experimental cost.

In this study, we combined antigen-binding B cell sorting with BCR-seq analysis to prioritize mABs with preferential isoD7-Aβ binding. Repertoires from isoD7-Aβ- and Aβ-binding B cell subsets were analyzed to prioritize IGH–IGK candidate combinations, which were then expressed and screened for specificity by ELISA and MST.

We defined clusters based on shared V-segment usage and CDR3 amino acid sequence similarity, aiming to group related clones that recognize similar epitopes and get useful signals of antigen specificity without requiring exact sequence matches. This approach can help identify sets of candidate antigen-associated clones across samples. To prioritize individual clonotypes and candidate heavy-light chain combinations, we extracted clonotype groups from clusters within specific samples and mice. We adopted a straightforward clustering approach complemented by somatic hypermutation (SHM) lineage tree analysis. However, the SHM tree analysis was less informative in our study because the experiment lacked multiple sequential timepoints and sufficient sequencing depth. The proposed workflow enabled the identification of IGH and IGK clonotypes resembling those previously reported to be specific for Aβ and/or isoD7-Aβ. In our analysis, we observed biased V-gene usage among IGH and IGK clonotypes in selected clusters, consistent with the stereotyped murine N-terminal anti-Aβ responses elicited by immunization, which frequently employ the IGHV8-12 together with IGKV1-117 [24]. In line with this, IGHV8-12 was used in the IGH clonotype #1263 of P4, the repertoire-derived variant that showed binding activity in both MST and ELISA assays. The IGH clonotypes #1263 and #1257, used in P4 and P4(Y104H), respectively, both belong to IGH cluster #48 and differ by only one amino acid in CDR3. Clonotype #1263 was detected as an IGHG1 clonotype but was represented by only a single UMI, whereas the closely related IGHM clonotype #1257 was more abundant in the cluster (Figure 2E).

Previous structural and alanine scanning studies of N-terminal anti-Aβ antibodies (PFA-1/2, WO2, 12A11, 10D5, 12B4) showed that recognition of the immunodominant Aβ_3–6_ epitope was driven primarily by germline-encoded residues in IGH CDR2, particularly the motif (W(Y)WDD(E)D), with additional contacts from IGK CDR1 and, to a lesser extent, IGH/IGK CDR3 [24, 27, 28]. Notably, those N-terminal antibodies commonly derive from IGHV8-12 and IGKV1-117 pairings.

However, the heavy–light chain pairings suggested by our workflow showed markedly different binding properties. Out of fourteen repertoire-derived mABs assessed in this study, four (P3, P4, P4(Y104H), and P13) showed binding in ELISA. Distinct binding profiles became apparent when full-length Aβ42 peptides were used for testing in ELISA. P3 bound both full-length isoforms weakly, P4 showed little binding, P4(Y104H) showed preferential binding to isoD7-Aβ42, and P13 displayed the clearest discrimination between isoD7-Aβ42 and Aβ42. This preference was further supported by the peptide-titration ELISA, in which P13 showed a concentration-dependent increase in binding to isoD7-Aβ42 while the signal for Aβ42 remained close to background (**Figure S10, Table S5**).

In subsequent MST affinity measurements, P4 bound isoD7-Aβ with an affinity approximately two orders of magnitude lower than that of K11, a previously published anti-isoD7-Aβ mAB (Kd = 1.6 ± 0.2 μM for P4 versus 18.9 ± 6.8 nM for K11). The dissociation constants of K11 measured by MST in this study differed by less than one order of magnitude from those measured by surface plasmon resonance in the original article [11].

Further highlighting convergence in our data, the light chain used in P4 and P4(Y104H) derives from the same V-gene segment, IGKV8-21, used by isoD7-Aβ-specific mAB K11, and differs from the K11 IGK by only four amino-acid substitutions. When this light chain was paired with the K11 heavy chain, the resulting hybrid antibody showed substantially higher affinity for isoD7-Aβ than when the same light chain was combined with the P4 heavy chain #1263 (IGHV8-12). These data suggest that IGHV8-12 provides the canonical germline-encoded scaffold for recognition of the N-terminal Aβ core (DAEFR), consistent with the conserved VH CDR2 motif characteristic of N-terminal anti-Aβ antibodies.

Our pairing results further suggest that IGKV8-21 may contribute to isoD7-Aβ recognition by shaping the heavy-light chain paratope geometry, although isoD7-Aβ selectivity is likely determined by the complete native heavy–light chain combination. IsoD7-Aβ differs from native Aβ by a subtle backbone isomerization at Asp7, which preserves the primary sequence while altering local geometry and probably epitope conformation. Consistent with a germline-favored N-terminal response, our repertoires were enriched for canonical IGK clonotypes (**Table S4**) which may retain cross-reactivity, whereas true isoD7 selectivity likely requires precise contacts at the Asp7-proximal edge of the epitope and the correct IGH-IGK chain paratope geometry.

The comparison of different heavy–light chain combinations supports this interpretation. The same IGHV10-1 heavy chain was tested in three combinations (P11–P13). No detectable binding was observed when it was paired with IGKV8-19 (P11) or IGKV1-117 (P12), whereas pairing with IGKV2-112 (P13) resulted in the clearest preferential recognition of isoD7-Aβ42. Notably, IGKV2- 112 was derived from a cluster composed exclusively of clonotypes derived from isoD7-Aβ-specific B-cell samples. This suggests that relatively rare light-chain clonotypes associated specifically with the antigen-binding compartment may contribute to discrimination between isoD7-Aβ and native Aβ. Taken together, our observations of V- and J-segment usage and sequence features similar to those reported in other studies suggest that mispairing is likely a major explanation for the limited binding activity observed and remains the principal challenge for further improvement of our current approach.

In this study, chain pairing was guided by visual inspection of heatmaps representing the frequencies of clonotype groups across corresponding replicates. While several approaches have been developed to automate TCR chain pairing (α and β chains) in high-throughput sequencing data, methods such as the Bayesian probability model MAD-HYPE [29] and its recent reimplementation TIRTL-seq [30] rely on Pearson correlation coefficients and require several dozen replicates to function optimally, which was not feasible for our dataset. In principle, there are no restrictions on applying a similar approach to BCRs. Though, in our setting the limited number of antigen-specific B cells makes such replicate-intensive approaches difficult to use for heavy-light chain pairing.

Our approach utilized tetramer sorting, which substantially reduces the labor intensity of screening and relies on bulk BCR-seq which is less costly than single-cell analysis [31]. Importantly, our method yielded BCR repertoires even when the number of sorted B cells was limited, allowing us to detect convergent candidate clonotypes under low-input conditions.

As a limitation of the study, our analysis revealed the relatively modest immunogenicity of the isoD7- Aβ_1-16_ conjugated to OVA and the overall low numbers of antigen-specific B cells. This also creates a practical balance between stringent sorting gate and the number of cells available for bulk repertoire analysis. While sorting on B cells that bind high amounts of antigen-containing tetramers or monomers is expected to increase the specificity and affinity of isolated B cells, low frequency of isoD7-Aβ-binding B cells (with an exception in the mouse # 6) necessitated the use of a less stringent sorting gate to recover a sufficient number of B cells for sequencing. Under these conditions, larger mouse cohorts and increased cell yields per sample may be required in such experiments to improve the detection of relevant clonotypes and the robustness of chain-pairing in bulk BCR-seq analyses.

Another limitation of our analysis is that clonotypes were grouped mainly by V/J usage and CDR3 amino-acid sequence similarity. This can merge related variants that differ in CDR1, CDR2, or framework regions and may therefore miss substitutions important for affinity. More detailed analysis of full-length variable-region sequences within selected clonotype groups may help identify improved variants for recombinant testing in future studies.

Overall, our results provide initial evidence that bioinformatics-guided repertoire analysis can prioritize antibody candidates, but also show clear gaps and limitations in the current workflow. A major source of false-positive candidates is the uncertainty of correct heavy–light chain pairing when repertoire analysis is performed on bulk or partially paired data. Incorrect pairing during recombinant mAB reconstitution can convert a true binder into a non-binder.

Importantly, the current study assesses antibody binding rather than biological activity. Affinity and preferential recognition of isoD7-Aβ are important criteria for further candidate selection, but the biological properties of the most promising mABs require additional validation and optimization. Although the results demonstrate that isoD7-dependent recognition can be retained in the full-length peptide, recognition of oligomeric or fibrillar isoD7-Aβ species was not examined. Epitope accessibility may differ in aggregated Aβ.

The present workflow offers a comparatively economical strategy for reducing an antigen-enriched repertoire to a small, experimentally tractable set of candidates. Although relatively few cells were obtained and analyzed here (a total of about 3.5 thousand isoD7-Aβ_1-16_-specific B cells from six mice), its throughput is essentially only limited by the number of animals available for immunisation. The “hit rate” observed here, roughly 4/14 (29%), was higher than that reported in some hybridoma-based studies, where a small fraction of screened clones turned out positive (*e.g.*, 8.7% in [32], but much lower than in some display screenings where hit rates could reach over 80% [33].

An advantage of our approach is that antibody candidate prioritization can be combined with analysis of the humoral response to the antigen of interest, as illustrated here by preferential V-gene usage in responses to Aβ and isoD7-Aβ. Despite the limitations associated with uncertain heavy–light chain pairing, bulk repertoire sequencing is technically simpler and less expensive than paired single-cell sequencing [31]. Further improvement of chain-pairing prediction may increase the efficiency of candidate selection while retaining these advantages.

## 4. Materials and Methods

### Mice and immunization

Six- to 7-week-old female BALB/c mice were obtained from the Center for Collective Use of the Institute of Physiologically Active Compounds and housed under SPF conditions at the Animal Facility of the Center for Precision Genome Editing and Genetic Technologies for Biomedicine, Engelhardt Institute of Molecular Biology, Russian Academy of Sciences (EIMB RAS). All animal procedures were performed in accordance with Russian regulations of animal protection and approved by the local Ethics Review Committee at EIMB RAS (Protocol No. 1 from 23/03/23).

To generate, identify, and isolate isoD7-Aβ_1-16_–specific B cells, six 6–7-week-old BALB/c mice were immunized intraperitoneally with ovalbumin-conjugated isoD7-Aβ_1-16_ (OVA–isoD7-Aβ_1-16_: H2N- DAFRH(isoD)SGYEVHHQK-OVA) formulated in Ribi adjuvant. All mice received a homologous booster with the same antigen 3 weeks after the prime (day 21). Two of the six mice received a second homologous booster 3 weeks later (day 42). Spleens were collected at three time points: 7 or 15 days after the first boost (n=2 per time point; days 28 and 36), and 7 days after the second boost (n=2; day 49).

### Staining and cell sorting

Splenocyte isolation, single cell preparation and staining with fluorescent antibodies and tetramers with isoD7-Aβ_1-16_ and Aβ – antigens was done as previously described [22]. Sorting was performed directly into 15 ul of UniSoRT Buffer (MiLaboratory) for subsequent cDNA synthesis to maximize recovery from low-input samples [34] B cells were directly sorted into lysis buffer optimized for downstream reverse transcription, thereby maximizing efficiency when working with low cell numbers (**Table S1**). Sorted B cell samples were frozen at −20C for storage prior to cDNA library preparation.

### cDNA library preparation

BCR repertoire cDNA libraries were prepared using the 5’RACE BCR (IGH/IGK) KIT MOUSE (MiLaboratory) reagent kit according to the manufacturer’s protocol (https://milaboratory.ru). Briefly, all sorted cells in the lysis buffer were used for the reverse transcription reaction with incorporation of UMIs via the template switch oligo. After first strand cDNA synthesis, the reaction was treated with uracil DNA glycosylase (UDG) to remove unincorporated UMI oligonucleotides, followed by purification using AMPure XP beads (Beckman Coulter). Purified cDNA was amplified in two consecutive PCR reactions according to the manufacturer’s protocol. The resulting IGH and IGL libraries were pooled. The concentration of pooled DNA amplicons was measured using Qbit 3 fluorometer. Fastseq300 platform was used to perform sequencing with 300 × 2 paired-end cycle kit. **Raw data treatment**

Raw sequencing reads were processed using MiXCR software (4.7) with “milab-mouse-rna-tcr-umi- race” preset. Processing included alignment to reference V(D)J libraries, Unique Molecular Identifiers (UMIs) extraction and correction, and clonotype assembly. Clonotype assemble procedure was performed on UMI-groups that passed the minimum read coverage threshold automatically selected by MiXCR (Table S2, overseq_threshold); UMI-groups with lower coverage were filtered out. On average, the sequencing coverage reached 45 reads per unique molecular identifier (UMI) for IGH and 17 reads per UMI for IGK, which is generally considered sufficient for downstream analysis (**Figure S1**, **Table S2**). The minimal thresholds applied had median values of 7 reads per UMI for IGH and 4 reads per UMI for IGL.

Allele variant calling was also performed using MiXCR 4.7 “findAlleles” subprogram [35], treating all mice as a single donor (we assumed that BALB/c inbred mice have the same allele variants). Somatic hypermutation (SHM) trees were constructed with MiXCR 4.7 “findShmTrees” subprogram with default settings. SHM tree construction was performed on all mouse IGH samples.

Immune repertoire statistics calculation and cluster analysis were performed using a custom “repseq” library (https://github.com/mmjmike/repseq). All clusters were created and analyzed using the “clustering” module from “repseq” library.

### IGH and IGK usage

IGH V-gene usage only for functional clonotypes was quantified from MiXCR output using UMI- weighted counts. For each sample, we computed the relative frequency of each IGHV segment as the fraction of UMIs assigned to clonotypes carrying that segment among all IGH clonotype UMIs in the sample. V-usage was then summarized at the group level by calculating the mean per-sample frequency for each IGHV segment within the iso-specific (N = 20), cross-specific (N = 13), and non- specific groups (N = 20). Statistical significance of the changes was estimated using one-sided t-test with subsequent FDR correction.

### Clustering analysis

Prior to clustering, each individual repertoire was subject to the “pooling” procedure: all clonotypes sharing the same combination of V/J-segments and CDR3 amino acid sequence (CDR3aa) were combined into a single one with the sum of UMI-counts and other non-additive features of the clonotype with the highest UMI-count in the group. All unique V-J-CDR3aa combinations (clonotype groups) throughout the whole dataset were given a consecutive numbering.

Clusters of IGH and IGK clonotypes were built based on the clonotype similarity: all functional clonotypes (defined as clonotypes without out-of-frame rearrangements or stop-codons in the CDR3 region) were included. Each clonotype was represented as a graph node. Nodes were connected by an edge if they shared the same V-segment and had at most one amino acid mismatch in the CDR3 region (ignoring the J-segment) – Hamming distance ≤1. Clusters themselves we define as the individual connected components of the graph, created as described above.

To select IGH clusters with isoD7-Aβ_1-16_-specific clonotypes, we used two filtering approaches; we retained clusters if one of the following conditions:

□ the cluster consists only of clonotypes with IGHV10-1 or IGHV14-3 segments, and contained at least two nodes derived from sorted isoD7-Aβ_1-16_-specific B cell samples,
□ the cluster contained nodes originated only from sorted isoD7-Aβ_1-16_-specific B cell samples and comprised at least 3 nodes

For selection of IGK clusters we used the following conditions:

□ the cluster consists only of clonotypes with IGKV1-110, IGKV1-117, or IGKV8-19 segments, and contained at least two nodes derived from sorted isoD7-Aβ_1-16_-specific B cell samples, or
□ the total UMI count of nodes from isoD7-Aβ_1-16_-specific samples was at least 100 in the cluster.

Additionally we filtered out clusters having cumulative UMI count <10 contributed by nodes from repertoires of sorted isoD7-Aβ_1-16_-specific B cells. From the selected clusters we have chosen the V-J- CDR3aa “pooled” clonotypes that were present in at least two isoD7-Aβ_1-16_-specific (or cross-specific) samples and had at least 10 total UMI count in these samples.

For selection of cross-specific clusters similar filters were applied.

### Visualization

Plots were created with the R language (“ggplot” and “pheatmap” libraries). Clusters were visualized using Cytoscape software. SHM trees visualization was performed with “ggtree” and “msa” R libraries.

Global embedding of IGH repertoires was generated by classical multidimensional scaling (MDS) based on pairwise sequence similarity. Two-dimensional embeddings of IGH repertoires (non-specific, iso-specific and cross-specific) generated from pairwise Hamming distances between aligned full-length nucleotide sequences. For each chain, all complete sequences were subjected to multiple sequence alignment using MAFFT v7.520 with default settings [36], after which the number of nucleotide differences was calculated for each pair of sequences without mismatch weighting. The resulting distance matrix was used for classical MDS to obtain a two-dimensional representation of repertoire structure. Each dot represents one clonotype/sequence, and dot size is proportional to UMI count.

### Recombinant antibody production in ExpiCHO-S suspension culture

DNA sequences encoding variable regions of light and heavy chains of K11 [11](European Patent Office EP3431496A1) and candidate light and heavy chains were acquired as fragments cloned into pUC19 vector (Cloning Facility). Plasmids were used to transform Escherichia coli cells, strain XL1- Blue (Evrogen) using Gene Pulser Xcell (Bio-Rad) according to the manufacturer’s protocol. Resulting bacterial colonies were grown in overnight cultures, and plasmid DNA was extracted from 5 ml of culture using Plasmid Miniprep kit (Evrogen) according to the manufacturer’s protocol. IGVH and IGVK sequences were cloned into pFUSEss-CHIg-hG4 and pFUSEss-CLIg-hK vectors (InvivoGen), respectively, via restriction-ligation using EcoRI and NheI (NEB) for IGH and EcoRI and PspLI (SibEnzyme) for IGK, and T4 DNA ligase (Promega). Plasmids contain sequences encoding constant regions of human IgG4 and IGK chains, therefore yielding chimeric mouse-human Ig genes.

Recombinant mABs were produced using the ExpiCHO-S suspension cell line (Thermo Fisher Scientific). Cells were cultured in 125-mL Erlenmeyer flasks containing 25 mL of ExpiCHO Expression Medium (Thermo Fisher Scientific) at 37°C. During routine cultivation, cells were counted every 3–4 days. The culture density was maintained between 0.1 × 10 and 6 × 10 cells/mL by replacing part of the culture with fresh medium.

For co-transfection, cells were grown to a density of 7–10 × 10 cells/mL, diluted to 6 × 10 cells/mL, and transfected with 0.5 µg of each plasmid encoding the heavy (HC) and light (LC) chains per mL of culture. Transfection was performed using OptiPRO SFM medium and the ExpiFectamine reagent (Thermo Fisher Scientific) according to the manufacturer’s instructions. Twenty-four hours after transfection, 0.006 volumes of ExpiCHO Enhancer and 0.24 volumes of ExpiCHO Feed (Thermo Fisher Scientific) were added to the culture.

After 8–10 days of cultivation, the medium containing secreted mABs was separated from cells and debris by centrifugation for 20 min at 4 °C and 350 × g, followed by 20 min at 4 °C and 2,200 × g. The supernatant was filtered through a 0.22-µm filter for subsequent affinity chromatography.

Purification of MAbs from culture medium was performed using 1 ml HiTrap Protein A HP columns (Cytiva) and a Bio-Lab 100 chromatography system (Hanbon Sci. & Tech.). The column was first equilibrated with 10 volumes of PBS, then culture medium was added at 1 ml/min flow rate. The column was washed with 10 volumes of PBS, then MAbs were eluted with an acidic buffer (0.1 M Gly-HCl, pH 2.8), collecting 500 µl fractions into tubes with 50 µl of neutralizing buffer (1 M Tris-HCl, pH 8.8). Columns were stored in 25% ethanol and reused up to 10 times.

Protein content in fractions was assessed by absorbance at 280 nm using NanoDrop One instrument (ThermoFisher). 2 µg of protein from three most protein-rich fractions were run on SDS-PAGE with 5% stacking gel, 10% separating gel with addition of trichloroethanol, using PageRuler Prestained Protein Ladder (ThermoFisher). Protein bands in gels were visualized using ChemiDoc XRS+ gel documentation system (Bio-Rad) using fluorescence of trichloroethanol (“stain-free” protocol). After ensuring that bands corresponded in mass to HC and LC, the three aforementioned fractions were pooled, transferred to PBS, and concentrated to a volume of approximately 100 μl in JetSpin 0.5 ml 10 kDa MWCO centrifugal concentrators (Jet Biofil). From 25 ml of cell culture (31 ml after addition of ExpiCHO Feed), approximately 1-1.5 mg of mABs were reproducibly obtained.

### ELISA

For ELISA, 50 μl of 0.1 mg/ml BSA-isoD7-Aβ_1-16_/BSA-Aβ_1-16_ (bovine serum albumin conjugated with isoD7-Aβ_1-16_ or Aβ_1-16_, degree of labeling 1:17, Peptide Specialty Laboratories) or 1 μM isoD7- Aβ_1-42_/Aβ1-42 (Peptide Specialty Laboratories) in bicarbonate buffer (2.33 g/l Na2CO3, 2.86 g/l NaHCO3, 0.2 g/l MgCl2, pH 9.8) were added to the wells of a 96-well plate and incubated for 1 h at room temperature.

Solutions were discarded with a sharp movement and wells were washed three times with 200 µl of PBST (0.05% Tween20), incubating each time for 5 min on an orbital shaker at 110 rpm and discarding liquid with a sharp movement. 300 µl of blocking buffer (PBST + 0.1% w/v fish gelatin (Biotium)) was added and the wells were incubated for 1 h at room temperature. The wells were washed three times as described above.

Then, 50 µl of mAB in PBS was added to the wells. Wells were incubated for 1 h at room temperature and washed three times as described above. 50 µl of secondary goat anti-human HRP-conjugated antibodies (ABClonal) (1:4000 dilution in PBS) were added to the wells. Wells were incubated for 45 min at room temperature and washed three times as described above. Then, wells were rinsed three times with PBS by adding 200 µl and discarding without incubation.

Finally, 100 µl of TMB solution (AbiZyme) was added to the wells. After color development, the reaction was stopped by adding 100 µl of 0.16 M H2SO4. TMB and H2SO4 were added to different wells at equal intervals, ensuring that all wells were incubated with TMB for the same amount of time. The signal was measured by absorbance at 450 nm using a plate reader. Each experimental point was performed in two technical replicates.

### Microscale Thermophoresis

The capacity of anti-isoD7-Aβ antibodies to bind isoD7-Aβ or Aβ were measured with the Microscale Thermophoresis (MST) [37]. Monomerization of Aβ peptides was made using hexafluoroisopropanol as described in [38] with following modifications. Immediately after addition of DMSO, stock solutions of the peptides were thoroughly vortexed for 30 sec and then ultra-sounded for 10 min at 20°C. K11, mK11, P3, and P4 mABs were stained with Alexa Fluor™ 488 Microscale Protein Labeling Kit (Thermo Fisher Scientific) according to the manufacturer’s protocol. Antibody concentration was kept constant at 2.5 nM while a final concentration of unlabeled Aβ ranged between

3.8 nM and 125 μM and a concentration of unlabeled isoD7-Aβ varied between 0.3 nM and 10 μM (in experiments with K11 and mK11) or between 3 nM and 100 μM (for binding with P4 and P5) with serial two-fold dilutions of the peptides. The assay buffer was PBST (Nano TemperTechnologies GmbH, München, Germany) containing 20% glycerol and 5% DMSO. Samples were loaded into Monolith NT.115 Premium Capillaries and MST analysis was performed using the Monolith NT.115 system (Nano Temper Technologies GmbH, München, Germany). Experiments were carried out in triplicate. Data analysis was performed using MO. Affinity Analysis software v.2.3.

Author Contributions: conceptualization: L.S.A., K.S.A., M.A.A., B.O.V., G.I.L., C.D.M.; methodology: G.T.V., B.O.V., G.I.L., M.E.M., K.E.A., K.O.I. and B.E.V.; software: M.M.Yu.; validation: M.E.M., N.N.A., K.E.A. and K.O.I.; formal analysis: M.M.Yu., K.E.A., K.O.I.; investigation: S.I.A., G.I.L., M.E.M., N.N.A., V.K.B., K.E.A., GTV and K.O.I.; resources: K.E.A., K.O.I., M.V.A., B.E.V.; data curation: M.M.Yu, K.E.A., K.O.I.; writing—original draft preparation: B.O.V., M.M.Yu., G.I.L., N.N.A., K.E.A. and K.O.I.; writing—review and editing: B.O.V., M.M.Yu., G.I.L. and N.N.A.; visualization: V.K.B., K.E.A., M.M.Yu. and N.N.A.; supervision, B.O.V., G.I.L., M.V.A., L.S.A. and M.A.A.; project administration: B.O.V., M.V.A, G.I.L.; funding acquisition: M.V.A. and M.A.A. All authors have read and agreed to the published version of the manuscript.

## Institutional Review Board Statement

The animal study protocol was approved by Russian regulations of animal protection and approved by the local Ethics Review Committee at EIMB RAS (Protocol No. 1 from 23/03/23).

## Data Availability Statement

The raw data supporting the conclusions of this/ article will be made available by the authors on request. The data are available in SRA: SUB15828023

## Acknowledgments

We acknowledge Irina Yu Petrushanko for technical assistance. Conflicts of Interest: The authors declare no conflicts of interest.

## Abbreviations

The following abbreviations are used in this manuscript:

mAB: Monoclonal antibody
BCR: B cell receptor
isoD7-Aβ_1-16_: Asp7-isomerized variant
OVA: Ovalbumin
CSW: Class-switched
IGH: Immunoglobulin heavy chain
IGK: Immunoglobulin kappa chain
MST: Microscale Thermophoresis
SHM: Somatic Hypermutation
FACS: Fluorescence Activated Cell Sorting
UMI MSA MDS: Unique Molecular Identifier Multiple Sequence Alignment Multidimensional Scaling

