## Supplementary material for "BCR repertoire analysis and cloning of antibody candidates targeting native and Asp7-isomerized β-amyloid": Supplemetary material

**Supplemental Material**

**Table S1. Number of sorted B cells using fluorescent tetramers with isoD7–Aβ_1-16_ or Aβ_1-16_.**

|  |  | **Specific to isoD7-Aβ** | | **Cross-specific to isoD7-Aβ and Aβ** | |
| --- | --- | --- | --- | --- | --- |
| **Mouse ID** | **Replica** | **memory isotype switched** | **GC B cells** | **memory isotype switched** | **GC B cells** |
| #4 (1) | R1 | 90 | 360 | - | 24 |
| #5 (3) | R1 | 100 | 200 | 65 | 200 |
|  | R2 | 55 | 40 | - | 131 |
| #6* (3) | R1 | 100 | 400 | 57 | 400 |
|  | R2 | 34 | 400 | - | 400 |
|  | R3 | - | 400 | - | - |
|  | R4 | - | 400 | - | - |
| #7 (1) | R1 | 127 | 140 | - | 192 |
|  | R2 | - | 67 | - | 93 |
| #8 (2) | R1 | 213 | 50 | 40 | 153 |
|  | R2 | - | - | - | - |
| #9 (2) | R1 | 86 | 207 | 8 | 100 |
|  | R2 | - | 83 | - | 52 |

#6* - plasmoblasts were sorted for this mouse

**Table S2. Quality control of sequencing data.**

| Sample_id | Chain | mouse_№ | Binder type | Cell count | B Cell type | Total reaads | Reads_aligned_pc | Total_umi | umi_after_correction | Overseq_threshold | Reads_after_filter | All clones | Reads_in_clones_total | Cones_func | Reads_in_func_clones |
| --- | --- | --- | --- | --- | --- | --- | --- | --- | --- | --- | --- | --- | --- | --- | --- |
| 4_Bcell_GC_cross_specific_2 | IGK | 4 | cross | 24 | GC | 3904 | 3,33 | 13 | 7 | 1 | 126 | 3 | 83 | 3 | 83 |
| 4_Bcell_GC_nonspecific_1 | IGH | 4 | non | 1000 | GC | 135019 | 60,12 | 5652 | 1791 | 7 | 79253 | 331 | 35743 | 294 | 32451 |
| 4_Bcell_GC_nonspecific_1 | IGK | 4 | non | 1000 | GC | 27926 | 79,85 | 1827 | 736 | 6 | 21414 | 331 | 18840 | 298 | 17472 |
| 4_Bcell_GC_nonspecific_2 | IGH | 4 | non | 1000 | GC | 104579 | 57,58 | 5167 | 2055 | 6 | 58639 | 309 | 25405 | 283 | 23827 |
| 4_Bcell_GC_nonspecific_2 | IGK | 4 | non | 1000 | GC | 23452 | 78,64 | 1640 | 762 | 6 | 17637 | 293 | 14428 | 265 | 13220 |
| 4_Bcell_GC_nonspecific_3 | IGH | 4 | non | 1000 | GC | 131531 | 64,39 | 6513 | 2050 | 7 | 82542 | 380 | 44115 | 344 | 40449 |
| 4_Bcell_GC_nonspecific_3 | IGK | 4 | non | 1000 | GC | 16490 | 83,06 | 1408 | 746 | 5 | 13037 | 280 | 10679 | 257 | 10035 |
| 4_Bcell_GC_nonspecific_4 | IGH | 4 | non | 1000 | GC | 80187 | 67,06 | 4914 | 1558 | 6 | 52368 | 247 | 23858 | 221 | 22259 |
| 4_Bcell_GC_nonspecific_4 | IGK | 4 | non | 1000 | GC | 26322 | 81,97 | 2163 | 758 | 6 | 20671 | 281 | 17554 | 264 | 16496 |
| 4_Bcell_GC_specific | IGH | 4 | iso | 360 | GC | 42946 | 54,25 | 1966 | 467 | 7 | 22856 | 70 | 9961 | 65 | 9507 |
| 4_Bcell_GC_specific | IGK | 4 | iso | 360 | GC | 4624 | 82,16 | 419 | 189 | 5 | 3644 | 54 | 2417 | 52 | 2352 |
| 4_Bcell_mem_isoAB_specific | IGH | 4 | iso | 90 | mem | 6612 | 35,16 | 267 | 51 | 11 | 2272 | 9 | 1270 | 9 | 1270 |
| 4_Bcell_mem_isoAB_specific | IGK | 4 | iso | 90 | mem | 513 | 82,26 | 39 | 15 | 1 | 415 | 7 | 372 | 7 | 372 |
| 5_Bcell_GC_cross_specific_1 | IGH | 5 | cross | 200 | GC | 21639 | 30,39 | 542 | 171 | 6 | 6366 | 46 | 3183 | 46 | 3183 |
| 5_Bcell_GC_cross_specific_1 | IGK | 5 | cross | 200 | GC | 1540 | 63,64 | 175 | 103 | 4 | 907 | 9 | 198 | 7 | 181 |
| 5_Bcell_GC_cross_specific_2 | IGH | 5 | cross | 131 | GC | 18789 | 49,26 | 550 | 200 | 7 | 8994 | 52 | 4051 | 51 | 3982 |
| 5_Bcell_GC_cross_specific_2 | IGK | 5 | cross | 131 | GC | 2133 | 74,4 | 223 | 154 | 4 | 1481 | 29 | 796 | 25 | 716 |
| 5_Bcell_GC_nonspecific_1 | IGH | 5 | non | 1000 | GC | 56214 | 71,43 | 3750 | 1468 | 5 | 39163 | 180 | 11944 | 168 | 11203 |
| 5_Bcell_GC_nonspecific_1 | IGK | 5 | non | 1000 | GC | 7132 | 79,15 | 1211 | 883 | 3 | 5181 | 140 | 2534 | 132 | 2387 |
| 5_Bcell_GC_nonspecific_2 | IGH | 5 | non | 1000 | GC | 54994 | 62,28 | 3678 | 1302 | 5 | 33369 | 115 | 6953 | 109 | 6639 |
| 5_Bcell_GC_nonspecific_2 | IGK | 5 | non | 1000 | GC | 8282 | 81,12 | 1338 | 1015 | 3 | 6164 | 194 | 3512 | 176 | 3219 |
| 5_Bcell_GC_nonspecific_3 | IGH | 5 | non | 1000 | GC | 52416 | 67,53 | 4332 | 1312 | 5 | 34376 | 108 | 7115 | 98 | 6539 |
| 5_Bcell_GC_nonspecific_3 | IGK | 5 | non | 1000 | GC | 7905 | 80,85 | 1415 | 1033 | 3 | 5851 | 143 | 2510 | 121 | 2207 |
| 5_Bcell_GC_specific_1 | IGH | 5 | iso | 200 | GC | 6324 | 26,83 | 195 | 116 | 4 | 1649 | 13 | 616 | 12 | 609 |
| 5_Bcell_GC_specific_1 | IGK | 5 | iso | 200 | GC | 1502 | 72,04 | 228 | 155 | 3 | 999 | 23 | 557 | 21 | 454 |
| 5_Bcell_GC_specific_2 | IGH | 5 | iso | 40 | GC | 5863 | 39,08 | 147 | 64 | 7 | 2230 | 7 | 1842 | 7 | 1842 |
| 5_Bcell_GC_specific_2 | IGK | 5 | iso | 40 | GC | 1107 | 42,73 | 80 | 47 | 4 | 450 | 4 | 155 | 4 | 155 |
| 5_Bcell_mem_isoAB_AB1_16_cross_specific | IGH | 5 | cross | 65 | mem | 6062 | 40,25 | 157 | 64 | 6 | 2378 | 9 | 821 | 8 | 803 |
| 5_Bcell_mem_isoAB_AB1_16_cross_specific | IGK | 5 | cross | 65 | mem | 318 | 73,27 | 48 | 39 | 3 | 202 | 5 | 84 | 3 | 65 |
| 5_Bcell_mem_isoAB_specific_1 | IGH | 5 | iso | 100 | mem | 13696 | 52,58 | 668 | 257 | 6 | 6993 | 45 | 3208 | 44 | 3163 |
| 5_Bcell_mem_isoAB_specific_1 | IGK | 5 | iso | 100 | mem | 3142 | 81,92 | 439 | 273 | 4 | 2384 | 47 | 1654 | 42 | 1577 |
| 5_Bcell_mem_isoAB_specific_2 | IGH | 5 | iso | 55 | mem | 11861 | 22,37 | 191 | 47 | 9 | 2593 | 16 | 1376 | 13 | 1179 |
| 5_Bcell_mem_isoAB_specific_2 | IGK | 5 | iso | 55 | mem | 772 | 60,49 | 67 | 50 | 4 | 451 | 11 | 273 | 11 | 273 |
| 6_Bcell_GC_cross_specific_1 | IGH | 6 | cross | 400 | GC | 44709 | 57,35 | 2065 | 765 | 6 | 24788 | 132 | 12898 | 127 | 12540 |
| 6_Bcell_GC_cross_specific_1 | IGK | 6 | cross | 400 | GC | 2945 | 64,89 | 407 | 315 | 3 | 1750 | 28 | 654 | 21 | 542 |
| 6_Bcell_GC_cross_specific_2 | IGH | 6 | cross | 400 | GC | 34994 | 65,62 | 1746 | 594 | 6 | 22153 | 115 | 13315 | 108 | 12721 |
| 6_Bcell_GC_cross_specific_2 | IGK | 6 | cross | 400 | GC | 1247 | 68,24 | 258 | 212 | 2 | 773 | 8 | 125 | 4 | 84 |
| 6_Bcell_GC_nonspecific_1 | IGH | 6 | non | 1000 | GC | 24730 | 66,85 | 1614 | 813 | 5 | 16109 | 56 | 2812 | 53 | 2672 |
| 6_Bcell_GC_nonspecific_1 | IGK | 6 | non | 1000 | GC | 7833 | 86,34 | 1145 | 804 | 3 | 6355 | 175 | 4212 | 152 | 3863 |
| 6_Bcell_GC_nonspecific_2 | IGH | 6 | non | 1000 | GC | 143691 | 59,41 | 5804 | 1958 | 7 | 83559 | 302 | 33026 | 269 | 31093 |
| 6_Bcell_GC_nonspecific_2 | IGK | 6 | non | 1000 | GC | 21401 | 78,38 | 2987 | 2293 | 3 | 15733 | 307 | 9288 | 253 | 8125 |
| 6_Bcell_GC_nonspecific_3 | IGH | 6 | non | 1000 | GC | 51711 | 60,63 | 2899 | 788 | 6 | 30527 | 93 | 7550 | 87 | 7137 |
| 6_Bcell_GC_nonspecific_3 | IGK | 6 | non | 1000 | GC | 5889 | 79,32 | 881 | 670 | 3 | 4306 | 116 | 2172 | 102 | 1974 |
| 6_Bcell_GC_specific_1 | IGH | 6 | iso | 400 | GC | 25100 | 55,64 | 1325 | 577 | 5 | 13498 | 76 | 6179 | 73 | 5955 |
| 6_Bcell_GC_specific_1 | IGK | 6 | iso | 400 | GC | 3508 | 78,91 | 586 | 432 | 3 | 2540 | 40 | 939 | 29 | 822 |
| 6_Bcell_GC_specific_2 | IGH | 6 | iso | 400 | GC | 34303 | 61,07 | 1740 | 555 | 6 | 20241 | 125 | 12011 | 116 | 11536 |
| 6_Bcell_GC_specific_2 | IGK | 6 | iso | 400 | GC | 2454 | 73,88 | 480 | 377 | 3 | 1555 | 26 | 426 | 25 | 421 |
| 6_Bcell_GC_specific_3 | IGH | 6 | iso | 400 | GC | 37328 | 61,23 | 1891 | 551 | 7 | 22090 | 129 | 12590 | 119 | 11808 |
| 6_Bcell_GC_specific_3 | IGK | 6 | iso | 400 | GC | 2651 | 69,63 | 380 | 299 | 3 | 1699 | 23 | 471 | 20 | 447 |
| 6_Bcell_GC_specific_4 | IGH | 6 | iso | 400 | GC | 22763 | 60,7 | 1137 | 460 | 6 | 13342 | 93 | 6132 | 86 | 5679 |
| 6_Bcell_GC_specific_4 | IGK | 6 | iso | 400 | GC | 2144 | 73,6 | 356 | 265 | 3 | 1425 | 27 | 458 | 25 | 447 |
| 6_Bcell_mem_isoAB_AB1_16_cross_specific | IGH | 6 | cross | 57 | mem | 4985 | 47,5 | 185 | 77 | 7 | 2285 | 16 | 1120 | 16 | 1120 |
| 6_Bcell_mem_isoAB_AB1_16_cross_specific | IGK | 6 | cross | 57 | mem | 240 | 65,83 | 42 | 31 | 3 | 142 | 1 | 28 | 1 | 28 |
| 6_Bcell_mem_isoAB_specific_1 | IGH | 6 | iso | 100 | mem | 14251 | 50,89 | 577 | 252 | 6 | 7031 | 35 | 4455 | 30 | 4296 |
| 6_Bcell_mem_isoAB_specific_1 | IGK | 6 | iso | 100 | mem | 4125 | 86,01 | 506 | 354 | 4 | 3335 | 21 | 2586 | 21 | 2586 |
| 6_Bcell_mem_isoAB_specific_2 | IGH | 6 | iso | 34 | mem | 3731 | 39,8 | 157 | 71 | 5 | 1455 | 1 | 32 | 1 | 32 |
| 6_Bcell_mem_isoAB_specific_2 | IGK | 6 | iso | 34 | mem | 591 | 75,8 | 81 | 54 | 4 | 414 | 8 | 223 | 7 | 207 |
| 6_Plasmobl_isoAB_AB1_16_cross_specific | IGH | 6 | cross | 467 | Plasm | 91770 | 71,88 | 6746 | 3502 | 4 | 63978 | 62 | 9153 | 58 | 9063 |
| 6_Plasmobl_isoAB_AB1_16_cross_specific | IGK | 6 | cross | 467 | Plasm | 12912 | 79,81 | 1934 | 1542 | 3 | 9441 | 17 | 1977 | 13 | 1935 |
| 6_Plasmobl_isoAB_specific | IGH | 6 | iso | 394 | Plasm | 82208 | 71,59 | 5538 | 1998 | 6 | 57396 | 86 | 14531 | 79 | 14095 |
| 6_Plasmobl_isoAB_specific | IGK | 6 | iso | 394 | Plasm | 15897 | 93,3 | 4030 | 3115 | 3 | 12720 | 36 | 5596 | 32 | 5561 |
| 7_Bcell_GC_cross_specific_2 | IGH | 7 | cross | 93 | GC | 47196 | 30,6 | 916 | 184 | 11 | 14022 | 55 | 6193 | 39 | 4557 |
| 7_Bcell_GC_cross_specific_2 | IGK | 7 | cross | 93 | GC | 1335 | 58,95 | 64 | 35 | 5 | 751 | 8 | 231 | 5 | 145 |
| 7_Bcell_GC_nonspecific_1 | IGH | 7 | non | 1000 | GC | 72909 | 49,49 | 2279 | 1047 | 6 | 35293 | 170 | 13135 | 151 | 12265 |
| 7_Bcell_GC_nonspecific_1 | IGK | 7 | non | 1000 | GC | 8515 | 78,44 | 742 | 485 | 4 | 6395 | 157 | 4294 | 134 | 3847 |
| 7_Bcell_GC_nonspecific_2 | IGH | 7 | non | 1000 | GC | 111677 | 37,36 | 2819 | 1206 | 6 | 40786 | 141 | 9258 | 122 | 8015 |
| 7_Bcell_GC_nonspecific_2 | IGK | 7 | non | 1000 | GC | 17316 | 75,88 | 1023 | 489 | 6 | 12588 | 202 | 9652 | 170 | 8336 |
| 7_Bcell_GC_nonspecific_3 | IGH | 7 | non | 1000 | GC | 73338 | 58,12 | 3113 | 944 | 7 | 41638 | 155 | 14476 | 143 | 13718 |
| 7_Bcell_GC_nonspecific_3 | IGK | 7 | non | 1000 | GC | 13571 | 84,55 | 1129 | 546 | 5 | 10983 | 153 | 7561 | 135 | 6941 |
| 7_Bcell_GC_nonspecific_4 | IGH | 7 | non | 1000 | GC | 97499 | 54,52 | 5439 | 1005 | 8 | 51564 | 150 | 17368 | 126 | 15241 |
| 7_Bcell_GC_nonspecific_4 | IGK | 7 | non | 1000 | GC | 12196 | 79,23 | 999 | 447 | 5 | 9159 | 187 | 7687 | 161 | 6774 |
| 7_Bcell_GC_specific_1 | IGH | 7 | iso | 140 | GC | 99743 | 25,81 | 1627 | 344 | 10 | 25256 | 71 | 14232 | 67 | 13796 |
| 7_Bcell_GC_specific_1 | IGL | 7 | iso | 140 | GC | 3043 | 62,44 | 177 | 92 | 8 | 1831 | 35 | 1423 | 0 | 0 |
| 7_Bcell_GC_specific_1 | IGK | 7 | iso | 140 | GC | 3043 | 62,44 | 177 | 92 | 8 | 1831 | 35 | 1423 | 32 | 1361 |
| 7_Bcell_GC_specific_2 | IGH | 7 | iso | 67 | GC | 18967 | 30,08 | 557 | 126 | 13 | 5515 | 23 | 3189 | 19 | 2785 |
| 7_Bcell_GC_specific_2 | IGK | 7 | iso | 67 | GC | 1412 | 65,3 | 84 | 35 | 12 | 889 | 13 | 666 | 13 | 666 |
| 7_Bcell_mem_isoAB_specific | IGH | 7 | iso | 127 | mem | 32829 | 42,64 | 871 | 165 | 20 | 13686 | 55 | 8988 | 53 | 8689 |
| 7_Bcell_mem_isoAB_specific | IGK | 7 | iso | 127 | mem | 1588 | 71,98 | 116 | 64 | 5 | 1095 | 22 | 894 | 20 | 854 |
| 8_Bcell_GC_cross_specific | IGH | 8 | cross | 153 | GC | 37391 | 49,53 | 1277 | 384 | 7 | 17966 | 127 | 12610 | 103 | 10459 |
| 8_Bcell_GC_cross_specific | IGK | 8 | cross | 153 | GC | 3189 | 73,47 | 263 | 119 | 5 | 2230 | 47 | 1572 | 32 | 1290 |
| 8_Bcell_GC_nonspecific_1 | IGH | 8 | non | 1000 | GC | 108203 | 53,92 | 4584 | 1166 | 7 | 57185 | 249 | 28157 | 228 | 26629 |
| 8_Bcell_GC_nonspecific_1 | IGK | 8 | non | 1000 | GC | 11930 | 77,21 | 1024 | 486 | 5 | 8730 | 207 | 7146 | 187 | 6370 |
| 8_Bcell_GC_nonspecific_2 | IGH | 8 | non | 1000 | GC | 82890 | 46,28 | 2523 | 710 | 8 | 37570 | 153 | 18164 | 146 | 17765 |
| 8_Bcell_GC_nonspecific_2 | IGK | 8 | non | 1000 | GC | 10112 | 69,15 | 535 | 276 | 6 | 6718 | 123 | 5359 | 112 | 4962 |
| 8_Bcell_GC_nonspecific_3 | IGH | 8 | non | 1000 | GC | 102123 | 44,18 | 3387 | 1152 | 7 | 44022 | 291 | 21966 | 271 | 20933 |
| 8_Bcell_GC_nonspecific_3 | IGK | 8 | non | 1000 | GC | 15390 | 78,06 | 1141 | 547 | 5 | 11503 | 218 | 9300 | 191 | 8276 |
| 8_Bcell_GC_specific | IGK | 8 | iso | 50 | GC | 1131 | 70,91 | 88 | 45 | 8 | 736 | 9 | 612 | 7 | 570 |
| 8_Bcell_mem_isoAB_AB1_16_cross_specific | IGH | 8 | cross | 40 | mem | 32021 | 13,43 | 1104 | 103 | 7 | 3891 | 4 | 432 | 4 | 432 |
| 8_Bcell_mem_isoAB_AB1_16_cross_specific | IGK | 8 | cross | 40 | mem | 659 | 38,09 | 31 | 15 | 1 | 243 | 6 | 172 | 6 | 172 |
| 8_Bcell_mem_isoAB_specific | IGH | 8 | iso | 213 | mem | 26990 | 52,11 | 1257 | 348 | 7 | 13678 | 76 | 7013 | 67 | 6367 |
| 8_Bcell_mem_isoAB_specific | IGK | 8 | iso | 213 | mem | 1305 | 75,79 | 156 | 88 | 4 | 919 | 29 | 756 | 28 | 743 |
| 9_Bcell_GC_cross_specific_1 | IGH | 9 | cross | 100 | GC | 54271 | 25,46 | 2329 | 256 | 9 | 12993 | 44 | 7329 | 43 | 7210 |
| 9_Bcell_GC_cross_specific_1 | IGK | 9 | cross | 100 | GC | 2058 | 62,78 | 145 | 71 | 4 | 1224 | 24 | 821 | 20 | 726 |
| 9_Bcell_GC_cross_specific_2 | IGH | 9 | cross | 52 | GC | 11957 | 41,11 | 306 | 93 | 9 | 4776 | 33 | 3424 | 32 | 3380 |
| 9_Bcell_GC_cross_specific_2 | IGK | 9 | cross | 52 | GC | 1208 | 67,55 | 91 | 40 | 11 | 779 | 14 | 521 | 11 | 433 |
| 9_Bcell_GC_nonspecific_1 | IGH | 9 | non | 1000 | GC | 156144 | 53,7 | 6224 | 1607 | 7 | 81898 | 213 | 33132 | 196 | 31724 |
| 9_Bcell_GC_nonspecific_1 | IGK | 9 | non | 1000 | GC | 17394 | 79,31 | 1396 | 537 | 6 | 13152 | 190 | 10490 | 171 | 9666 |
| 9_Bcell_GC_nonspecific_2 | IGH | 9 | non | 1000 | GC | 133224 | 54,2 | 4986 | 1391 | 7 | 70605 | 300 | 32771 | 277 | 31421 |
| 9_Bcell_GC_nonspecific_2 | IGK | 9 | non | 1000 | GC | 24329 | 82,5 | 1502 | 601 | 6 | 19185 | 262 | 16453 | 239 | 14969 |
| 9_Bcell_GC_nonspecific_3 | IGH | 9 | non | 1000 | GC | 127539 | 59,93 | 5976 | 1603 | 7 | 74607 | 318 | 34612 | 290 | 32274 |
| 9_Bcell_GC_nonspecific_3 | IGK | 9 | non | 1000 | GC | 21715 | 81,03 | 1397 | 562 | 6 | 16866 | 241 | 14477 | 222 | 13524 |
| 9_Bcell_GC_specific_1 | IGH | 9 | iso | 207 | GC | 20249 | 18,02 | 228 | 84 | 10 | 3548 | 22 | 1975 | 22 | 1975 |
| 9_Bcell_GC_specific_1 | IGK | 9 | iso | 207 | GC | 2696 | 50,26 | 137 | 66 | 6 | 1300 | 22 | 694 | 22 | 694 |
| 9_Bcell_GC_specific_2 | IGH | 9 | iso | 83 | GC | 4985 | 22,77 | 47 | 26 | 17 | 1125 | 4 | 424 | 4 | 424 |
| 9_Bcell_GC_specific_2 | IGK | 9 | iso | 83 | GC | 506 | 61,26 | 33 | 18 | 1 | 300 | 5 | 167 | 5 | 167 |
| 9_Bcell_mem_isoAB_AB1_16_cross_specific | IGH | 9 | cross | 8 | mem | 3348 | 10,87 | 23 | 5 | 1 | 348 | 2 | 239 | 2 | 239 |
| 9_Bcell_mem_isoAB_specific | IGH | 9 | iso | 86 | mem | 17962 | 33,41 | 476 | 73 | 19 | 5847 | 25 | 3724 | 20 | 3294 |
| 9_Bcell_mem_isoAB_specific | IGK | 9 | iso | 86 | mem | 554 | 84,66 | 52 | 26 | 7 | 448 | 9 | 425 | 9 | 425 |

**Table S3. Description of the IGH and IGK isoD7-Aβ specific and cross-specific clonotypes**.

|  | Clonotype ID | Cluster ID | Specificity | Chain | IGV | IGJ | CDR3 | Mice | MiXCR tree ID | Isotypes |
| --- | --- | --- | --- | --- | --- | --- | --- | --- | --- | --- |
| 1 | IGH_2735 | IGH_4 | isoD7-Aβ | IGH | IGHV14-3 | IGHJ2 | CASHYYGRSLFDYW | 9 |  | IGHG1 |
| 2 | IGH_1664 | IGH_11 | isoD7-Aβ | IGH | IGHV3-1 | IGHJ2 | CARRDGNGFDYW | 6 | 170 | IGHD, IGHM |
| 3 | IGH_2639 | IGH_55 | isoD7-Aβ | IGH | IGHV1-69 | IGHJ1 | CARYGYYLYFDVW | 6 |  | IGHD, IGHG1, IGHM |
| 4 | IGH_541 | IGH_84 | isoD7-Aβ | IGH | IGHV1-75 | IGHJ2 | CAREDYYGDFDYW | 6 | 2 | IGHG1, IGHM |
| 5 | IGH_664 | IGH_57 | isoD7-Aβ | IGH | IGHV2-6-8 | IGHJ2 | CAREPPGPYYFDYW | 6 |  | IGHG1, IGHM |
| 6 | IGH_1949 | IGH_12 | isoD7-Aβ | IGH | IGHV8-12 | IGHJ4 | CARRQLGLDAMDYW | 6 |  | IGHM |
| 7 | IGH_2344 | IGH_30 | isoD7-Aβ | IGH | IGHV8-12 | IGHJ4 | CARSRLRDYAMDYW | 6 |  | IGHG1, IGHM |
| 8 | IGH_1257/1263 | IGH_48 | isoD7-Aβ | IGH | IGHV8-12 | IGHJ4 | CARIYHYGSSAMDYW/CARIYYYGSSAMDYW | 6 |  | IGHM/IGHG1 |
| 9 | IGH_3612 | IGH_1 | isoD7-Aβ | IGH | IGHV10-1 | IGHJ2 | CVRDLFDYW | 4,6,7 | 60 | IGHG1, IGHM |
| 10 | IGH_3609 | IGH_1 | isoD7-Aβ | IGH | IGHV10-1 | IGHJ2 | CVRDLFDNW | 4,7 |  | IGHG1 |
| 11 | IGH_1132 | IGH_47 | isoD7-Aβ | IGH | IGHV9-3 | IGHJ2 | CARGYWFDYW | 6 |  | IGHM |
| 12 | IGH_3578 | IGH_18 | isoD7-Aβ | IGH | IGHV14-3 | IGHJ2 | CVDYDGYFW | 7 |  | IGHM |
| 13 | IGH_3598 | IGH_149 | isoD7-Aβ | IGH | IGHV10-1 | IGHJ3 | CVNDYGKGYW | 7 | 61 | IGHG1 |
| 14 | IGKL_3968 | IGK_21 | isoD7-Aβ | IGK | IGKV8-21 | IGKJ1 | CKQSYNLRTF | 6,8 |  | IGKC |
| 15 | IGKL_5185 | IGK_7 | isoD7-Aβ | IGK | IGKV1-110 | IGKJ5 | CSQSTHVPLTF | 4,5,6,7,8,9 |  | IGKC |
| 16 | IGKL_3843 | IGK_1 | isoD7-Aβ | IGK | IGKV1-117 | IGKJ1 | CFQGSLVPWTF | 9 |  | IGKC |
| 17 | IGKL_3839 | IGK_1 | isoD7-Aβ | IGK | IGKV1-117 | IGKJ1 | CFQGSHVPWTF | 4,6,9 |  | IGKC |
| 18 | IGKL_3846 | IGK_1 | isoD7-Aβ | IGK | IGKV1-117 | IGKJ1 | CFQGSRVPWTF | 9 |  | IGKC |
| 19 | IGKL_3834 | IGK_1 | isoD7-Aβ | IGK | IGKV1-117 | IGKJ5 | CFQGSHVPLTF | 6,7 |  | IGKC |
| 20 | IGKL_3840 | IGK_1 | isoD7-Aβ | IGK | IGKV1-117 | IGKJ2 | CFQGSHVPYTF | 6,7,8 |  | IGKC |
| 21 | IGKL_4956 | IGK_0 | isoD7-Aβ | IGK | IGKV6-15 | IGKJ5 | CQQYNSYPLTF | 4,5,6,7 |  | IGKC |
| 22 | IGKL_4955 | IGK_0 | isoD7-Aβ | IGK | IGKV6-15 | IGKJ4 | CQQYNSYPLTF | 6,8 |  | IGKC |
| 23 | IGKL_4953 | IGK_0 | isoD7-Aβ | IGK | IGKV6-15 | IGKJ1 | CQQYNSYPLTF | 5,7 |  | IGKC |
| 24 | IGKL_4334 | IGK_6 | isoD7-Aβ | IGK | IGKV8-19 | IGKJ2 | CQNDHSYPYTF | 5,6,7 |  | IGKC |
| 25 | IGKL_4345 | IGK_6 | isoD7-Aβ | IGK | IGKV8-19 | IGKJ5 | CQNDYSYPLTF | 4,5,6,7,8,9 |  | IGKC |
| 26 | IGKL_4909 | IGK_107 | isoD7-Aβ | IGK | IGKV4-54 | IGKJ5 | CQQYHSYPLTF | 5,6 |  | IGKC |
| 27 | IGH_1244 | IGH_7 | cross | IGH | IGHV8-12 | IGHJ4 | CARISRPAGAMDYW | 9 | 127,128,38 | IGHG2B, IGHG2C, IGHG3 |
| 28 | IGH_3415 | IGH_0 | cross | IGH | IGHV8-12 | IGHJ4 | CTRRGLYYGNFDAMDYW | 5 |  | IGHG1, IGHG2C |
| 29 | IGH_1374 | IGH_178 | cross | IGH | IGHV8-12 | IGHJ3 | CARLYYGNYW | 6 |  | IGHG1 |
| 30 | IGH_1853 | IGH_42 | cross | IGH | IGHV8-12 | IGHJ2 | CARRGYGFDYW | 6 |  | IGHM |
| 31 | IGH_2221 | IGH_66 | cross | IGH | IGHV8-12 | IGHJ1 | CARSGYFDVW | 6 |  | IGHG1, IGHM |
| 32 | IGH_1413 | IGH_274 | cross | IGH | IGHV8-12 | IGHJ1 | CARNGNFDVW | 6 |  | IGHE, IGHM |
| 33 | IGH_1586 | IGH_537 | cross | IGH | IGHV8-12 | IGHJ4 | CARRAEMDYW | 5,6 |  | IGHG1, IGHM |
| 34 | IGH_2070 | IGH_131 | cross | IGH | IGHV8-12 | IGHJ4 | CARRVTLITTVKGTDAMDYW | 6 |  | IGHG1, IGHM |
| 35 | IGH_1993 | IGH_63 | cross | IGH | IGHV8-12 | IGHJ4 | CARRSLRDEDAMDYW | 6 |  | IGHG1 |
| 36 | IGH_2385 | IGH_320 | cross | IGH | IGHV8-12 | IGHJ3 | CARSWDGYDEFAYW | 6 |  | IGHM |
| 37 | IGH_2676 | IGH_99 | cross | IGH | IGHV8-12 | IGHJ2 | CARYYGSSYGYW | 6 |  | IGHG1, IGHM |
| 38 | IGH_3095 | IGH_689 | cross | IGH | IGHV8-12 | IGHJ4 | CSRRALNYYGSNYDAMDYW | 6 |  | IGHM |
| 39 | IGH_1232 | IGH_8 | cross | IGH | IGHV8-8 | IGHJ4 | CARIPHYVSGAMDQW | 5 | 113 | IGHG1 |
| 40 | IGH_1804 | IGH_76 | cross | IGH | IGHV8-12 | IGHJ3 | CARRGLYDYDWFDYW | 9 | 20 | IGHG1 |
| 41 | IGKL_5202 | IGK_7 | cross | IGK | IGKV1-110 | IGKJ5 | CSQTTHVPLTF | 5,9 |  | IGKC |
| 42 | IGKL_5185 | IGK_7 | cross | IGK | IGKV1-110 | IGKJ5 | CSQSTHVPLTF | 5,6,8,9 |  | IGKC |
| 43 | IGKL_3834 | IGK_1 | cross | IGK | IGKV1-117 | IGKJ5 | CFQGSHVPLTF | 5,6,7,8,9 |  | IGKC |
| 44 | IGKL_3845 | IGK_1 | cross | IGK | IGKV1-117 | IGKJ5 | CFQGSRVPLTF | 5,8,9 |  | IGKC |

**Table S4. Characteristics of the top IGK clusters by isoD7-Aβ specific or cross-specific UMI count (count ≥ 100 or both count ≥ 25 and specific UMI proportion = 100 %).**

| Cluster ID | Specificity | specific UMI count | Nodes total | Total UMI count | specific UMI proportion | Mice | Consensus CDR3 sequence | V segment | Most frequent J segment |
| --- | --- | --- | --- | --- | --- | --- | --- | --- | --- |
| IGK_21 | isoD7-Aβ | 437 | 39 | 551 | 79 % | 6,8 | CKQSYNLRTF | IGKV8-21 | IGKJ1 |
| IGK_6 | isoD7-Aβ | 211 | 84 | 332 | 64 % | 4,5,6,7,8,9 | CQNDYSYPLTF | IGKV8-19 | IGKJ5 |
| IGK_0 | isoD7-Aβ | 155 | 194 | 1030 | 15 % | 4,5,6,7,8 | CQQYNSYPLTF | IGKV6-15 | IGKJ2 |
| IGK_1 | isoD7-Aβ | 148 | 129 | 476 | 31 % | 4,5,6,7,8,9 | CFQGSHVPLTF | IGKV1-117 | IGKJ5 |
| IGK_107 | isoD7-Aβ | 137 | 2 | 137 | 100 % | 5,6 | CQQYHSYPLTF | IGKV4-54 | IGKJ5 |
| IGK_7 | isoD7-Aβ | 101 | 89 | 381 | 26 % | 4,5,6,7,8,9 | CSQSTHVPLTF | IGKV1-110 | IGKJ1 |
| IGK_55 | isoD7-Aβ | 28 | 14 | 28 | 100 % | 5,6,7 | CQQLVEYPFTF | IGKV2-112 | IGKJ4 |
| IGK_1 | cross | 194 | 129 | 476 | 41 % | 5,6,7,8,9 | CFQGSHVPLTF | IGKV1-117 | IGKJ5 |
| IGK_7 | cross | 205 | 89 | 381 | 54% | 5,6,8,9 | CSQSTHVPLTF | IGKV1-110 | IGKJ1 |

**Table S5. Results of binding assays performed for most prominent candidate mABs, compared to K11. Where possible, K_D_ values are listed; otherwise, binding is specified qualitatively in relation to that of K11 with isoD7-Aβ in the same assay, denoted here as “+++”; “++” – weaker binding, “+” – much weaker binding, “-“ – no binding.**

| Antibody | IGHV gene | IGKV gene | ELISA isoD7-Aβ_1-16_ | ELISA Aβ_1-16_ | ELISA isoD7-Aβ_42_ | ELISA Aβ_42_ | ELISA isoD7-Aβ_42_, varying Aβ conc. | ELISA Aβ_42_, varying Aβ conc. | MST isoD7-Aβ_42_ | MST Aβ_42_ |
| --- | --- | --- | --- | --- | --- | --- | --- | --- | --- | --- |
| P3 | IGHV1-75 | IGKV8-19 | + | + | + | + | + | + | - | - |
| P4 | IGHV8-12 | IGKV8-21 | + | + | - | - | + | + | 1.6 μM | - |
| P4(Y104H) | IGHV8-12 | IGKV8-21 | + | + | + | - | + | + | - | - |
| P13 | IGHV10-1 | IGKV2-112 | ++ | + | ++ | - | 2.9 μM | - | - | - |
| K11 | IGHV2-9 | IGKV8-21 | +++ | +++ | +++ | ++ | 1.9 nM | + | 18.9 nМ | ⩾ 20 μM |

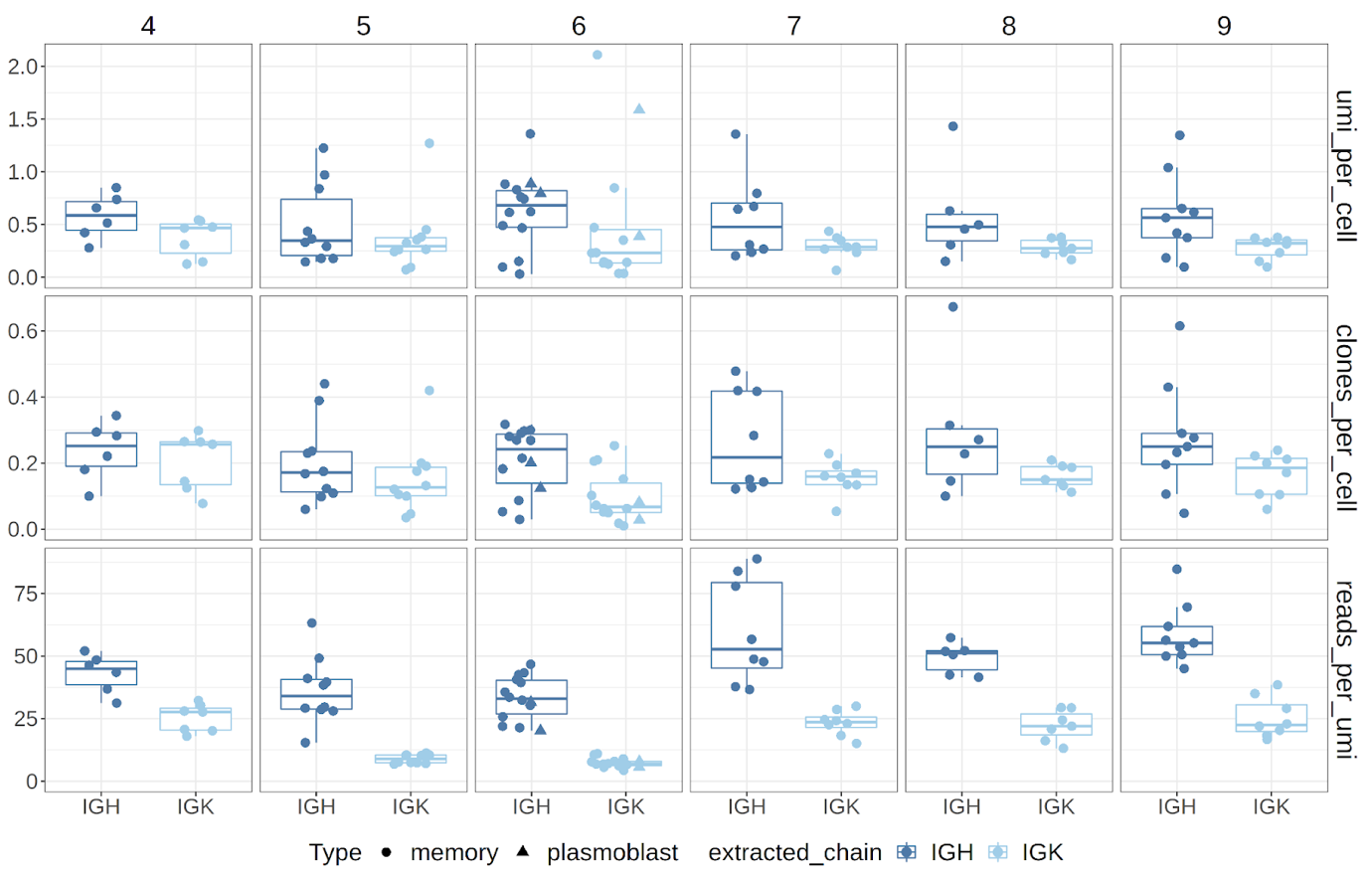

**Figure S1. Quality control of the IGH and IGK sequencing data.** Boxplots summarizing key quality control metrics for heavy (IGH) and light (IGK) chain BCR sequencing from sorted memory B cells (circles) and plasmablasts (triangles) across six mouse samples (IDs 4–9). Rows indicate three quality metrics: UMI per cell (top), number of clones per cell (middle), and reads per UMI (bottom). IGH and IGK data are shown separately for each mouse, colored dark and light blue, respectively. This visualization highlights per-sample consistency, technical variance between IGH and IGK, and confirms successful BCR reconstruction.

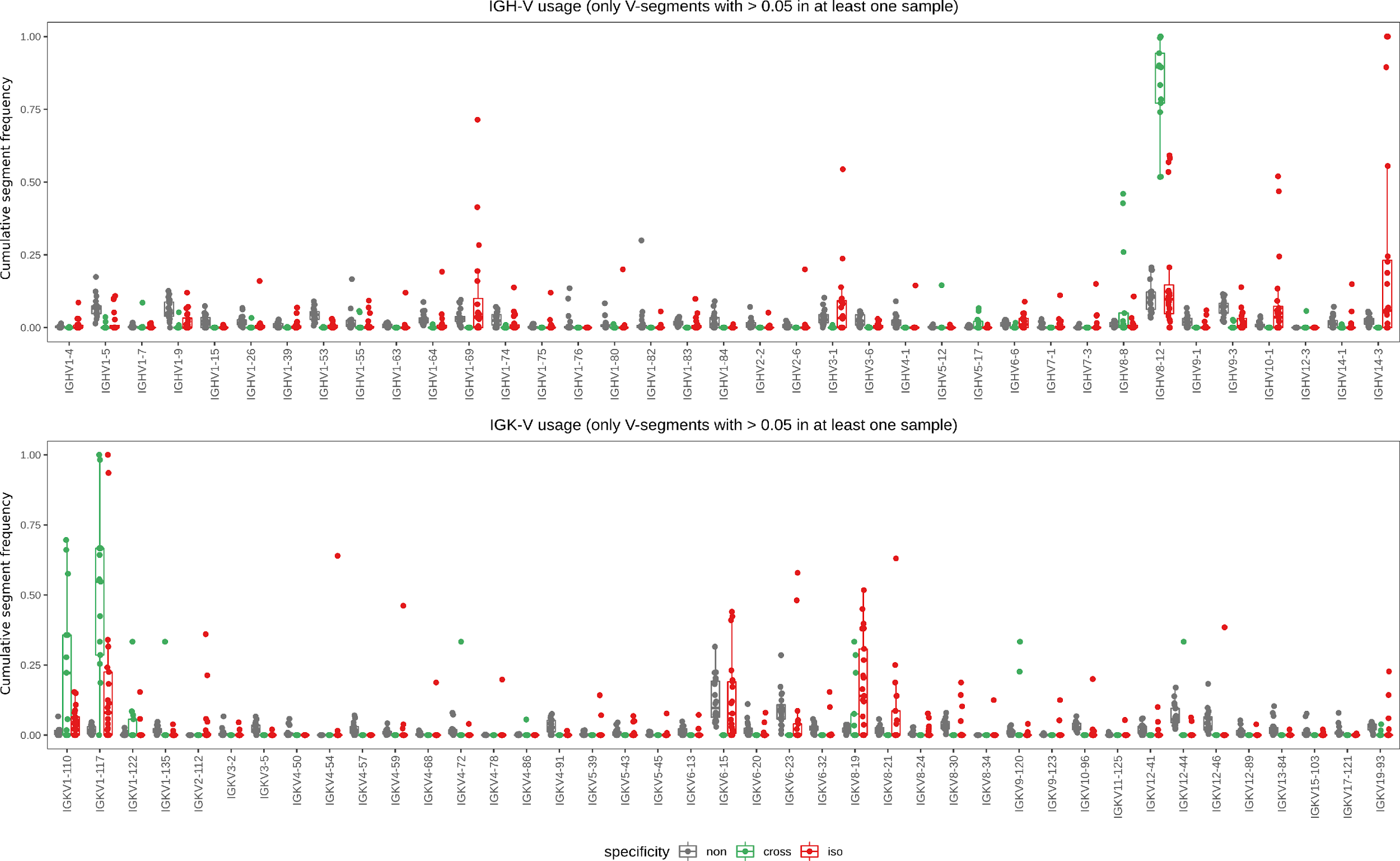

**Figure S2.** Cumulative V-segment frequency by UMI counts: IGH (upper) and IGK (lower). Each dot represents a sample. Only shown are the segments having a frequency higher than 0.05 in at least one sample. Samples are grouped and colored by specificity type: non-specific (N = 20 for both IGH and IGK), cross-specific (N = 13 for both IGH and IGK) and isoD7-Aβ_1-16_-specific (N = 20 for IGH and N = 21 for IGK).

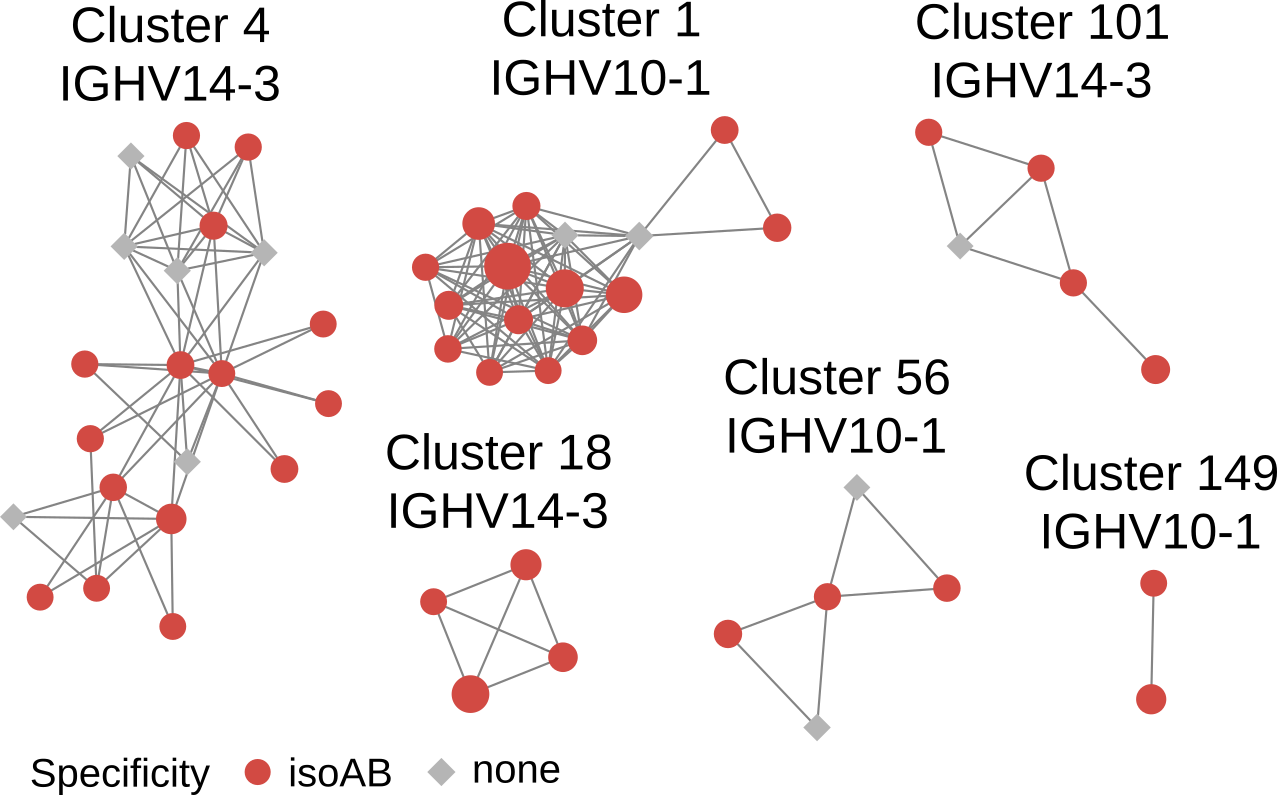

**Figure S3**. IGH Clusters, which were selected by the presence of significantly more represented V-segments (IGHV10-1 and IGHV14-3) in isoD7-Aβ-specific samples. Clonotypes from isoD7-Aβ_1-16_ sorted B cells are shown as red circles, and the ones from non-specific repertoires – as grey diamonds. The size of each node is proportional to its UMI count. Shown are the V-segments shared by each node in the cluster.

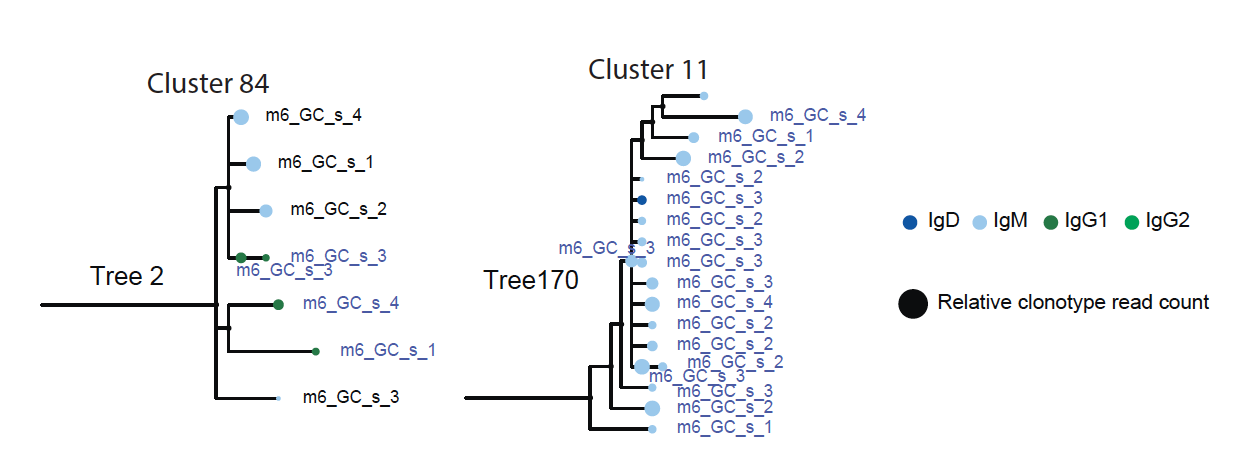
**Figure S4.** Somatic hypermutation (SHM) lineage trees reconstructed using MiXCR v4.7 from selected IGH clonotype groups derived from isoD7–Aβ-specific (s), cross-reactive (c), or non-specific (n) repertoires. Each leaf is labeled with the sample ID (e.g., m4_GC_s_1 = mouse #4, germinal center, specific), with node size indicating relative read count. Isotypes (IgD, IgM, IgG1, IgG2) are shown with color-coded dots.

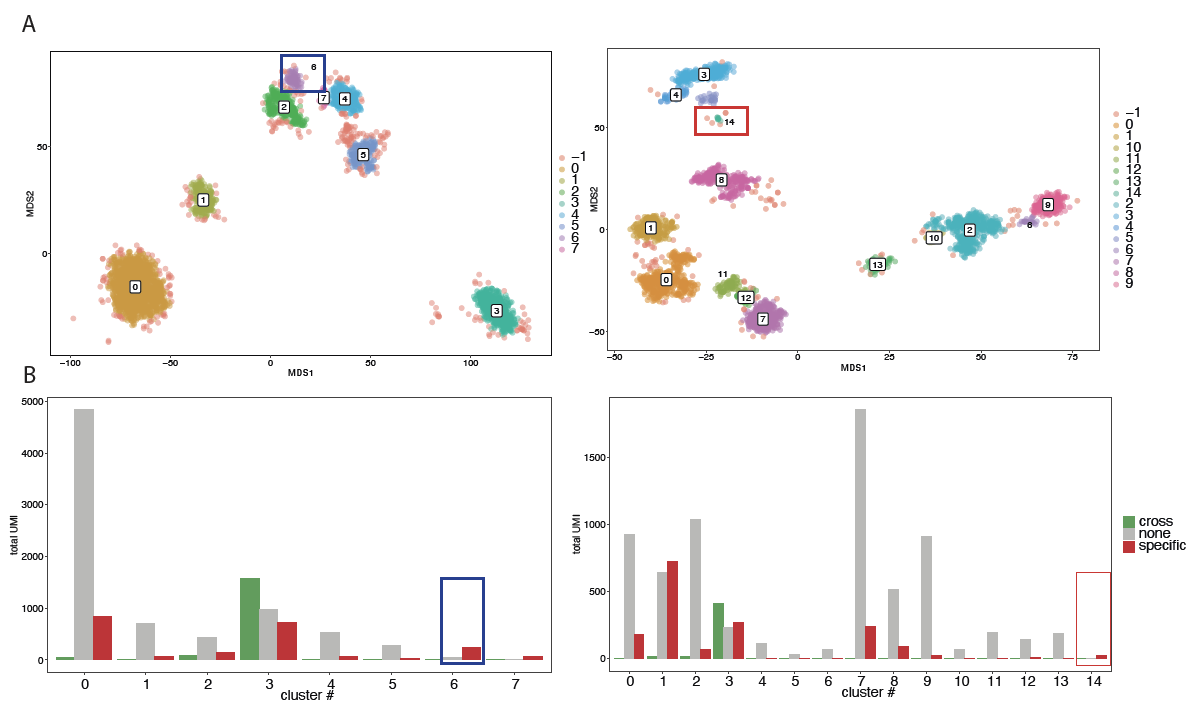

**Figure S5.** (A) The global embeddings of IGH and IGK repertoires generated by multidimensional scaling (MDS) based on pairwise Hamming distances between MAFFT-aligned full-length nucleotide sequences. Each dot represents one clonotype, and dot size is proportional to its UMI count in the repertoire. Dots are colored by DBSCAN clusters (IGH clustering hyperparameters: eps=2.5, min_samples=20; IGK clustering hyperparameters: eps=1.75, min_samples=8). (B) Respective total UMI counts by specificity groups are shown for each DBSCAN cluster. The outlined regions indicate DBSCAN clusters #6 for IGHV10-1 (blue) and #14 for IGKV2-112 (red), which showed enrichment for clonotypes derived from isoD7-Aβ1–16-specific repertoires.

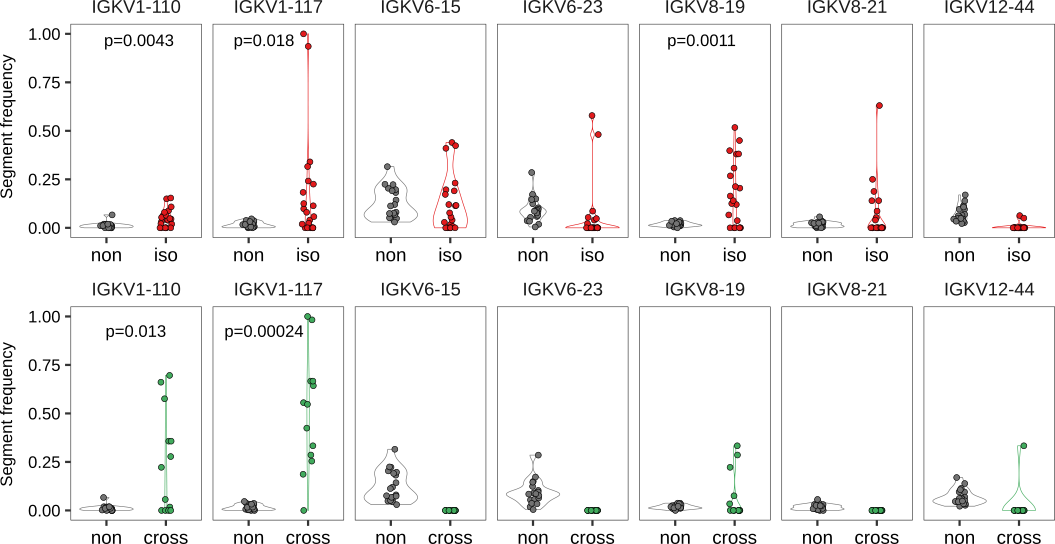

**Figure S6.** The top seven IGKV segments were selected based on a mean frequency threshold (>0.025) across all samples. Comparison groups were IGKV abundance between repertoires of non-specific (N = 20) and isoD7-Aβ_1-16_-specific (N = 21) repertoires and between repertoires from non-specific (N = 20) and cross-specific (N = 13) sorted B cells. P values were obtained using a one-sided t-test and adjusted for multiple comparisons using the Benjamini–Hochberg false discovery rate (FDR) correction.

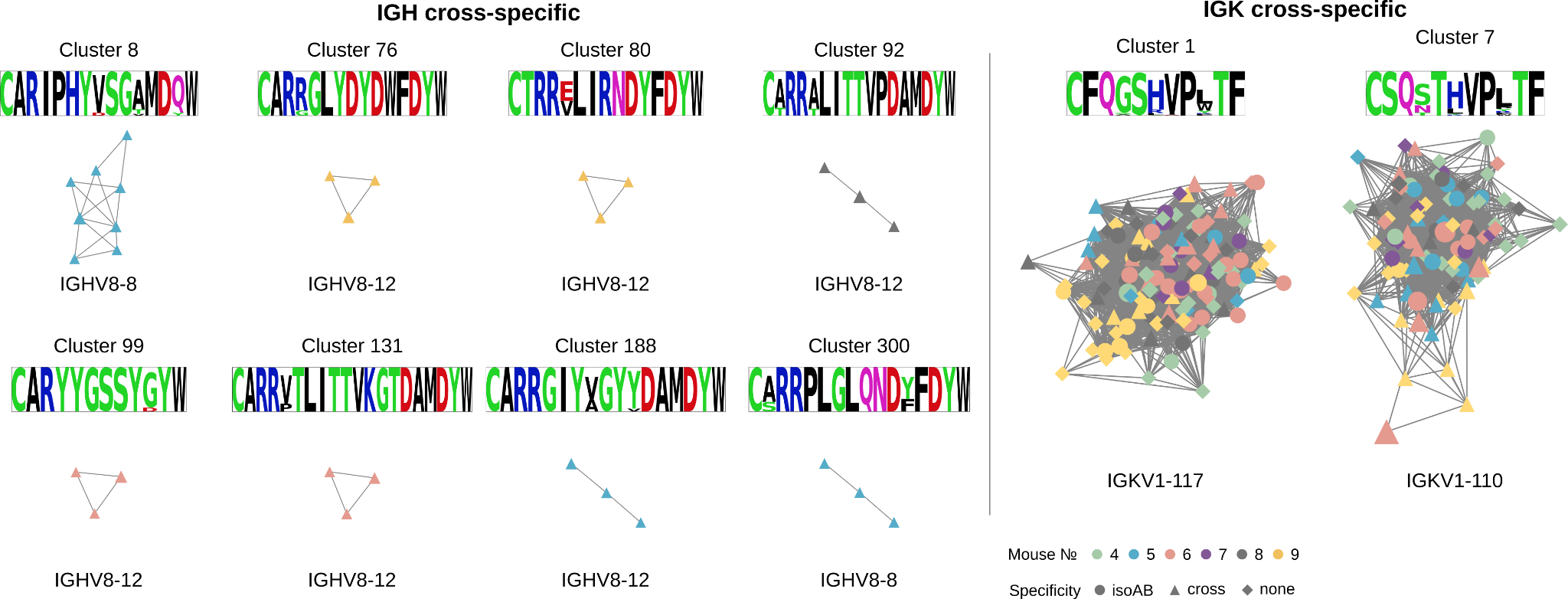

**Figure S7.** Representative examples of IGH and IGK clonotype clusters in network graphs built from the repertoires of sorted B cell populations grouping clonotypes that shared the same IGHV gene and differed by no more than one amino acid in the CDR3 region (≤1 aa mismatch). Shown IGH clusters (left panel) met the following criteria: contained only nodes originated from sorted cross-specific B cell samples (≥3 nodes) and cumulative UMI count contributed by nodes was ≥10. Shown IGK clusters (right panel) with cumulative UMI counts ≥100 of the cross-specific sample-derived nodes. Colors indicate the mouse of origin (m4–m9). CDR3 logos (weighted by UMI counts) of the clonotypes comprising each cluster are shown as well as V-segments which are shared between the nodes.

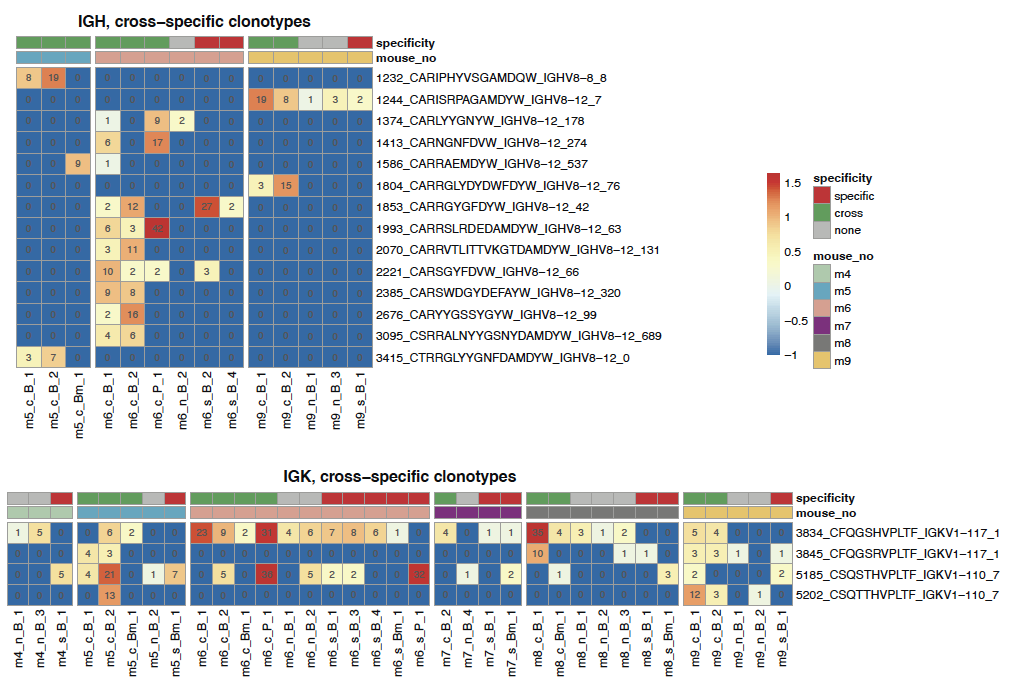

**Figure S8.** Heatmaps showing the abundance of selected IGH and IGK cross-specific clonotype groups across samples, expressed as log10 UMI counts. Samples are ordered by specificity (iso-specific, cross-specific, or non-specific) and by mouse ID. UMI counts in the sample are shown in each box. The annotations on the right show the clonotype ID, CDR3 amino acid sequence, V- and J-segments, and cluster ID in which the clonotypes were found.

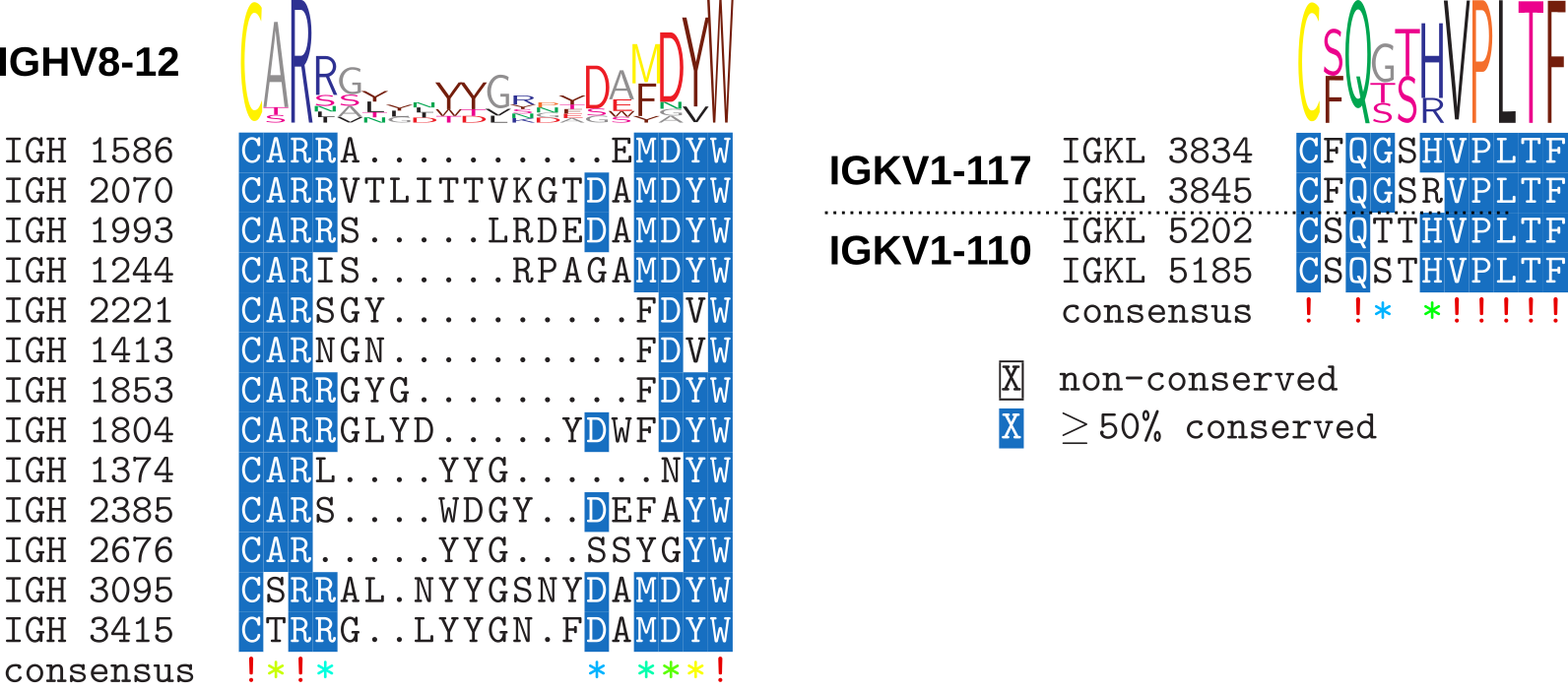

**Figure S9.** Multiple sequence alignment of the CDR3 regions of the selected cross-specific clonotypes. For IGH alignment, only one clonotype (IGH_1232) was excluded as it was the only one containing the IGHV8-8 segment. IGK CDR3 sequences for cross-specific clonotypes demonstrate high convergence, while IGH CDR3 sequences are diverse and mostly contain “sticky” residues like Gly and Tyr in the center of the CDR3 segments.

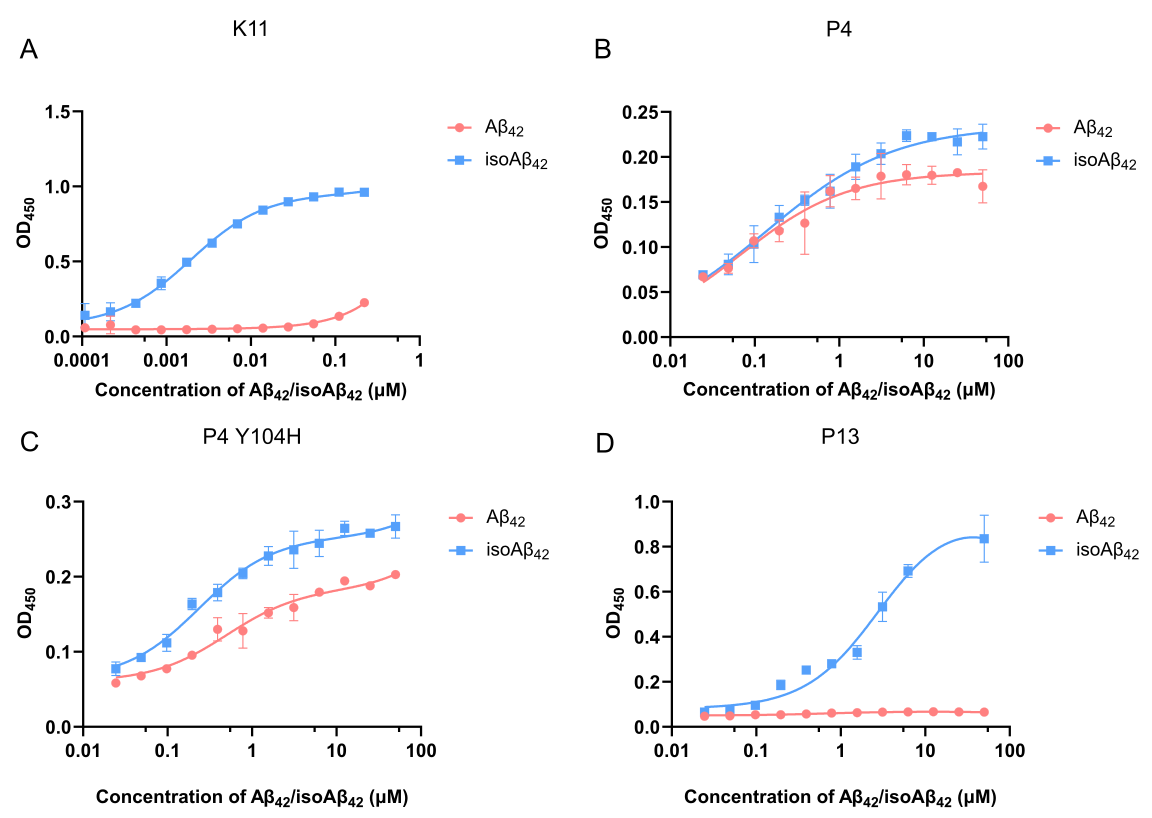

**Figure S10.** Binding of antibodies K11 (A), P4 (B), P4(Y104H) (C), and P13 (D) to in ELISA with varying Aβ_42_ or isoD7-Aβ_42_ peptide concentrations. The dependence of optical density (450 nm) on the concentration of Aβ_42_ and isoAβ_42_ (µM) is shown. Mean values ± standard deviations (SD) are presented (n=3).

**Supplemental Methods**

**ELISA with varying Aβ_42_/isoD7-Aβ_42_ concentrations**

Serial dilutions of synthetic Aβ_42_ and isoAβ_42_ peptides were prepared in PBS, ranging from 0.1 nM to 0.2 µM (for K11 antibody) or from 25 nM to 50 µM (for P4, P4(Y104H), and P13 antibodies). The peptides were added in triplicate to the wells of a 96-well ELISA plate (NEST) and incubated overnight at +4°C. The wells were washed three times with PBST (0.05% Tween 20) and blocked with 1% BSA (Dia-M) in PBST for 1 hour at room temperature with agitation. The wells were washed twice with PBST, and antibodies K11, P4, P4(Y104H), and P13 were added at a concentration of 2 µg/mL in PBST. After a 2-hour incubation at room temperature with agitation, the wells were washed three times with PBST and HRP-conjugated anti-human antibodies (HyTest) were added at a concentration of 0.4 µg/mL, followed by incubation for 30 minutes at room temperature. The wells were washed six times with PBST, and the signal was developed using TMB (Dia-M). The reaction was stopped by adding 0.25 M sulfuric acid, and the optical density was measured at 450 nm.
